# Deconstructing a behavioral state: parallel neural integrators control distinct features of an aversive behavioral state in *C. elegans*

**DOI:** 10.64898/2026.06.04.729906

**Authors:** Saba N. Baskoylu, Amit Kundra, Sarah Pugliese, Jungsoo Kim, Kamal Maher, Alex Hiser, Frank Meng, Cassi Estrem, Di Kang, Eric Bueno, Steven W. Flavell

## Abstract

Transient sensory stimuli can trigger long-lasting changes in behavior and cognition. For example, repeated aversive experiences can lead to persistent changes in arousal and mood. Specialized integrator circuits can store information about recent experiences, but how they are embedded within the architecture of the brain to control internal brain states remains poorly understood. We find that *C. elegans* accumulates evidence of past aversive experiences over minutes to generate a scalable behavioral state consisting of multiple behavioral changes. Using a brain-wide imaging screen, we identified a set of neurons that integrate aversive sensory stimuli over different timescales, from seconds to minutes. Neuronal perturbations show that these integrator neurons are critical for the aversive behavioral state. They control distinct behavioral features, such as changes in locomotion speed or increased responsiveness to subsequent aversive stimuli. Moreover, the integrator neurons function in parallel to one another, utilize different mechanisms to generate persistent neural activity, and signal through different transmitters. Our result show how a behavioral state can be decomposed into distinct behavioral features that map onto different neural integrators that act in parallel. Combined action of the integrators then generates the full set of behaviors that comprise the global state.

## Introduction

Animals must store information about their sensory surroundings to adapt their behavior to their environment. Neural integrators are neurons or circuits that stably integrate their input stimuli, allowing them to exhibit persistent neural responses to transient stimuli. They are thought to allow animals to build internal models of their external environments, for example by storing spatial information or modulating global brain states in response to environmental cues. However, how neural integrator circuits are embedded within the overall architecture of nervous systems to control brain states remains poorly understood.

Targeted studies of neural integrators have suggested potential mechanisms that may underlie their function as they accumulate evidence to control internal states. For example, network-level interactions have been implicated in stable attractor dynamics in the anterior lateral motor cortex (ALM) of mice during a short-term memory task^1^. P1 command neurons induce persistent activity in downstream pCD neurons to maintain a state of social arousal during Drosophila courtship^2^. Cell-intrinsic mechanisms for persistent neural activity during sensory integration have also been documented. Persistent accumulation of intracellular cAMP levels underlies evidence accumulation in mice and flies^3,4^. Neuromodulatory receptor activation has been implicated in generating persistent activity, for example by contributing to stable line attractor dynamics in the hypothalamus that control a state of aggressiveness^5^. These studies suggest that both neural circuit and cell-intrinsic mechanisms can contribute to persistent activity and neural integrator function, perhaps functioning together^6^. Identifying mechanisms of persistent activity across entire nervous systems and relating this to global brain states remains an ongoing challenge for the field.

Here, we use the simple model system *C. elegans* to ask how recent sensory experiences are stably represented in the brain to control an aversive behavioral state. With the readily available wiring diagram^7–9^, brain-wide imaging^10–13^, and multi-color atlas for cell identification^14^, it is feasible to link sensory experience and behavior to brain-wide dynamics^15^ in *C. elegans*. Because the same neural population can be examined across animals, this can be combined with precise genetic tools to dissect the function of individual neurons and circuits. This approach can link large-scale neural activity in behaving animals to fine-grained neural mechanisms.

*C. elegans* exhibits many persistent behavioral states. For example, foraging animals switch between minutes-long roaming and dwelling states while exploring bacterial food patches^16–19^. In addition, animals exhibit prolonged quiescence states during development and in response to stress, infection, and satiety^20–24^. Transient sensory stimuli can also have long-lasting effects on *C. elegans* behavior. A pair of nociceptive sensory neurons, called ASH neurons, detect aversive mechanical and chemical stimuli and drive an immediate escape response^25–27^. The escape response consists of a reversal and a turn, which are controlled by a defined network of interneurons^28–33^. However, repeated activation of the ASH nociceptive neurons in foraging animals leads to a period of elevated locomotion that lasts for minutes^34,35^. Repeated cross-modal activation of other aversive pathways has also been shown to elevate locomotion through a mechanism that requires RID neurons and FLP-3 neuropeptides^36^. It remains unclear how repeating sensory stimuli are integrated across the *C. elegans* nervous system, for example which neurons across the brain exhibit persistent activity and how this gives rise to a change in behavioral state.

In this study, we examined the brain-wide basis of neural integration in the context of aversive sensory stimulation. We determine that animals integrate their detection of aversive sensory cues over minutes, accumulating sensory information to control multiple behavioral parameters that define an aversive behavioral state: increased speed, reduced feeding rates, increased sensory responsiveness, and more. We then examine whole-brain neural dynamics in freely-moving animals, identifying specific neurons in the *C. elegans* nervous system that act as neural integrators, stably accumulating sensory information. Two integrator neurons with minutes-long persistent activity function in parallel to control distinct features of this aversive state: the neuron AVH stably integrates aversive stimuli to control speed, while the neuron PVQ integrates aversive stimuli to induce a priming effect, increasing the animal’s responsiveness to future aversive stimuli. We further show that AVH and PVQ rely on different neural mechanisms to generate persistent activity and utilize different transmitters to signal to downstream circuits. Our findings suggest an organization where multiple integrators can process sensory cues in parallel to induce distinct behavioral changes, together giving rise to the behavioral changes that comprise a global state.

## Results

### Repeated activation of the nociceptive sensory neuron ASH over minutes alters multiple motor programs, inducing an aversive behavioral state

To study temporal evidence accumulation in *C. elegans* circuits, we examined how repeated stimulation of the nociceptive sensory neuron ASH impacts the animal’s behavior. We chose to study ASH because it is the sensory neuron that is most reliably activated by aversive stimuli^37,38^. Acute ASH activation is known to induce reversals, whereas repeated activation results in fast forward movement^25,34^. To precisely control the timing of aversive cues and deliver them as programmable sequences, we began by optogenetically activating ASH (see below for experiments using natural stimuli). We identified the *nlp-76* promoter as driving strong, highly specific expression in ASH only (Extended Data Figure 1a) and used it to express the light-activated opsin CoChR. We delivered 2-second light pulses every 10 seconds to freely-moving animals on uniform bacterial lawns (Fig. 1a). Consistent with prior work^34^, repeated ASH photoactivation reliably triggered reversals at stimulus onset and progressively increased forward velocity during the inter-stimulus interval and after the last stimulus (Fig.1b). Repeated ASH stimulation also modulated other behaviors, gradually reducing both feeding and defecation rates (Fig. 1c-d). These behavioral changes, especially the speeding, persisted for several minutes after the final ASH stimulus (Fig. 1b-d), suggesting that repeated activation of ASH leads to a long-lasting change in multiple behavioral parameters.

**Figure 1.**
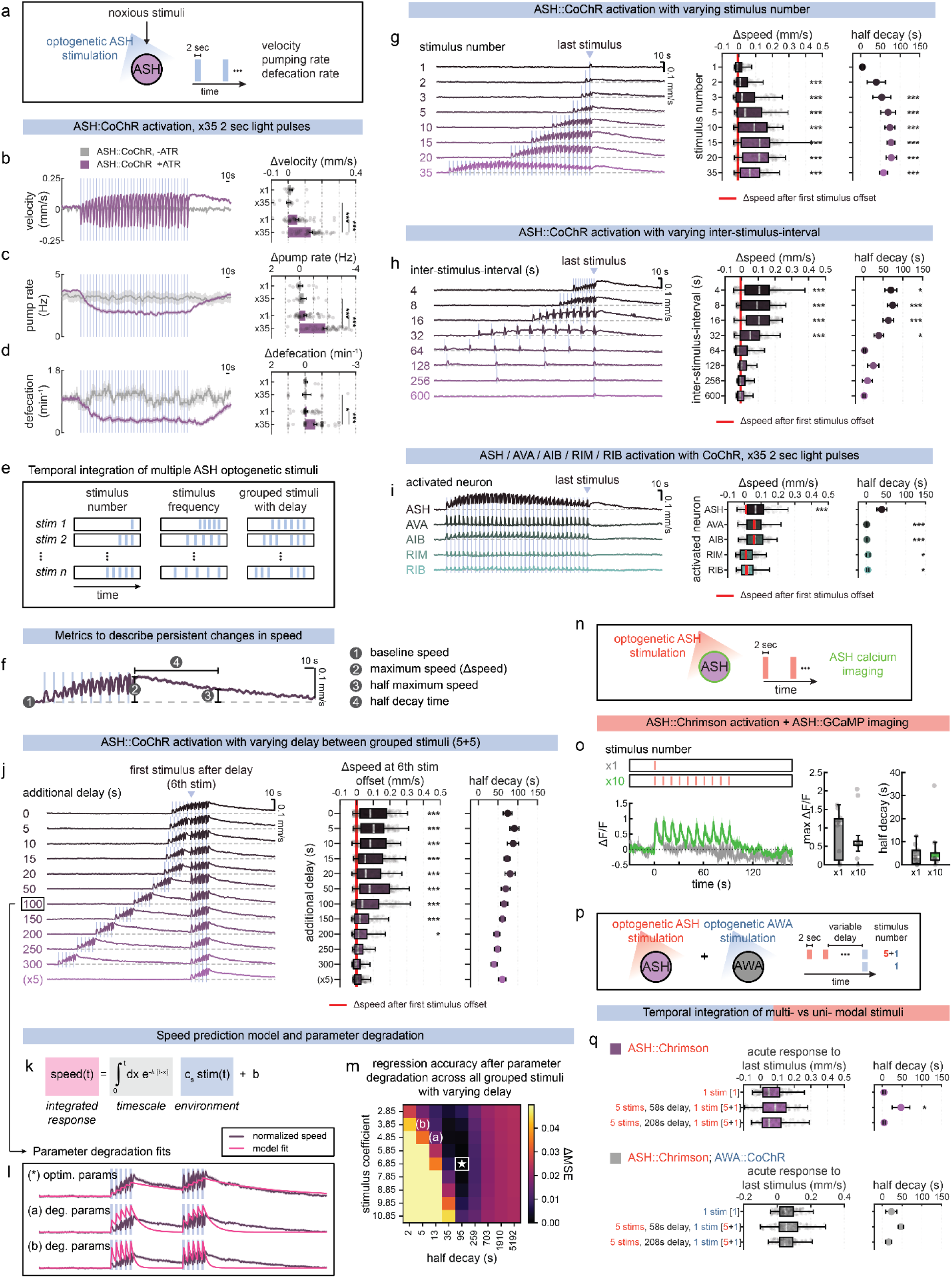
Repeated activation of ASH nociceptive neurons over minutes induces an aversive behavioral state consisting of multiple behavioral changes. a. Schematic of optogenetic ASH neuron activation and simultaneous behavioral recordings in freely-moving animals. Noxious stimuli (harsh touch, heavy metals, aversive odors) endogenously activate ASH neurons. To mimic this activation, animals expressing CoChR in ASH received repeated 2-second light pulses; velocity, pumping, and defecation frequency were assessed. b. Left: Event-triggered changes in velocity after repeated optogenetic ASH neuron activation. Right: Changes in speed after single vs. repeated stimuli. Error bars show mean ± SEM. Each behavior compared to no-ATR controls lacking the essential opsin co-factor all trans-retinal (ATR) that is supplied to *C. elegans* in the growth plate. ***p<0.001, Wilcoxon rank-sum test. n=62 animals for +ATR, n=15 animals for -ATR. c. Left: Event-triggered changes in pumping rate after repeated optogenetic ASH neuron activation. Right: Changes in speed after single vs. repeated stimuli. Error bars show mean ± SEM. Each behavior compared to no-ATR controls. ***p<0.001, Wilcoxon rank-sum test. n=62 animals for +ATR, n=15 animals for -ATR. d. Left: Event-triggered changes in defecation after repeated optogenetic ASH neuron activation. Right: Changes in speed after single vs. repeated stimuli. Error bars show mean ± SEM. Each behavior compared to no-ATR controls. *p<0.05, ***p<0.001, Wilcoxon rank-sum test. n=62 animals for +ATR, n=15 animals for -ATR. e. Schematic of different optogenetic ASH neuron activation patterns. Variable number: 2-second stimuli delivered 1, 2, 3, 5, 10, 15, 20, or 35 times at fixed 8-second intervals. Variable-frequency: 2-second stimuli delivered with 4, 8, 16, 32, 64, 128, 256, or 600 second inter-stimulus-intervals for a total of 10 times. Variable-delay: groups of 5 2-second stimuli delivered with various delays (8 - 308 seconds). f. Quantification of behavioral changes after repetitive ASH stimulation. Shown: average speed trace during optogenetic ASH neuron activation with 8-second inter-stimulus-intervals for 10 times. Gray dashed line indicates baseline speed. Four metrics of the behavioral trace are indicated. g. Repeated ASH optogenetic activation for variable number of times. Left: event-triggered speed changes. Middle: average speed change at offset of last stimulus. Right: half-decay time of average speed at stimulus offset as it regresses to baseline. Äspeed box plot: ***p<0.001, Bonferroni-corrected empirical p-value for speed changes at stimulation offset vs. first stimulus offset. Red dot and line show median and min-max whiskers for the first stimulus offset response pooled across all conditions. Boxes show the median and interquartile range (25th-75th percentile). Whiskers extend to the most extreme value within 1.5x the interquartile range. Dots are individual animals. Half-decay plot: ***p<0.001, Bonferroni-corrected empirical p-value for comparison to 1 stimulus condition. Error bars show standard deviation across bootstrap resamples. n = 3 recorded plates with 45-78 animals per plate. h. Repeated ASH optogenetic activation with varying inter-stimulus interval, shown as in (g). Half-decay plot: *p<0.05, ***p<0.001, Bonferroni-corrected empirical p-value for comparison to longest inter-stimulus-interval (600 seconds). n = 2-3 recorded plates with 48-104 animals per plate. i. Speed changes after repeated optogenetic activation of ASH neurons or command interneurons, shown as in (g). Left: event-triggered speed changes after repeated activation of indicated neuron at 8-second inter-stimulus-intervals for 35 times. Middle: average speed change at stimulus offset. Right: half-decay time of average speed at stimulus offset to baseline. Δspeed plot: ***p<0.001, Bonferroni-corrected empirical p-value for speed changes at stimulation offset vs. first stimulus offset. Half-decay plot: *p<0.05, ***p<0.001, Bonferroni-corrected empirical p-value comparing to ASH neurons. n = 2-4 recorded plates with 21-55 animals per plate. j. Event-triggered speed changes in response to two groups of 5 2-sec stimuli, with the groups separated by variable delays, shown as in (g). Middle: average speed change after first stimulus following delay (i.e. 6^th^ total stimulus). Right: half-decay time of average speed after last stimulus as it regresses to baseline. Last row: control with 5 stimuli alone. Δspeed plot: *p<0.05, ***p<0.001, Bonferroni-corrected empirical p-value for speed change at stimulation offset vs first stimulus offset. n = 4-8 recorded plates with 24-83 animals per plate. k. Mathematical model where animal speed is modeled as a function of optogenetic stimuli. Speed is predicted based on integrating recent stimuli with exponential decay. This model was trained on datasets with 4-, 16-, and 64-second inter-stimulus-interval ASH activation; and validated and tested on 32-second and 8-second inter-stimulus-interval datasets, respectively. t: time; speed[t]: predicted animal speed; integral: accumulated responses from time 0 to t; λ: decay rate; cs: stimulus coefficient; stim[x]: input signal (optogenetic stimuli) at past time x; b: baseline speed. l. Representative parameter degradation test for a single stimulus condition (100s delay). Shown: actual normalized speed (dark purple) and fits (pink) for optimal decay and stimulus coefficient parameters, plus fits after two parameter degradations (a and b). m. Model accuracy for parameter degradation, varying stimulus coefficient and decay parameter. Star indicates the trained model parameters. Optimal parameter combination with the lowest error is indicated in white box. Heatmap shows change in mean squared error (MSE) across all stimulus conditions relative to fit with trained parameters. Example degradation parameters (from 1l) are marked as (a) and (b). n. Illustration of optogenetics and calcium recording experiments in ASH neurons. Animals expressing Chrimson and GCaMP in ASH neurons received 2-second light pulses; changes in ASH calcium dynamics were assessed. o. Optogenetic ASH neuron activation with simultaneous ASH calcium recordings. Left: event-triggered averages of normalized ASH calcium dynamics after single or repeated ASH::Chrimson stimulation. Middle: maximum change after last stimulus offset. Right: half-decay time of maximum change at last stimulus offset. Boxes show the median and interquartile range (25th-75th percentile). Whiskers extend to the most extreme value within 1.5x the interquartile range. Dots are individual animals. n=11 animals for x1; n=10 animals for x10. p. Schematic of dual optogenetic setup. Left: ASH and AWA neurons express red light-activated Chrimson and blue-light activated CoChR, respectively. Right: ASH was repeatedly activated with red light (2-second stimuli delivered at 8-second inter-stimulus-interval 5 times), and after a short (58s) or long (208s) delay, AWA was activated once with blue light (2-second stimulus). Responses were compared to controls receiving a single AWA stimulus with no prior optogenetic stimulation. q. Repeated optogenetic ASH activation with red light-activated Chrimson, followed by either a sixth ASH stimulus with red light (top) or a single AWA neuron activation with blue light-activated CoChR (bottom) after either a short or long time lag. Left: magnitude of behavioral speeding responses to the last stimulus, measured from the 8-second baseline period preceding it. Right: persistence of behavioral response to the last stimulus as speed decays to baseline. All conditions compared to controls receiving a single AWA or ASH stimulus with no prior optogenetic stimulation. * p<0.05, empirical p-value compared to single-stimulus controls. For ASH::Chrimson, n = 2-4 recorded plates with 19-62 animals per plate. For ASH::Chrimson; AWA::CoChR, n = 3-4 recorded plates with 20-67 animals per plate.

We next varied the optogenetic stimulation pattern to assess how ASH stimuli are integrated to affect behavior (Fig. 1e). To capture the natural variation in aversive landscapes, we systematically varied the number (overall hazard level), frequency (temporal patterning), temporal grouping (clustered versus distributed hazards), intensity, and frequency trajectory (deteriorating, stable, or improving conditions) of ASH photoactivation (Fig. 1e, Extended Data Figure 1b-c). We then examined behavioral responses, quantifying the overall speed increase at stimulus offset, and the decay rate as speed regressed to baseline (Fig. 1f; speed was chosen as a behavioral readout here since it can be easily recorded in large populations of animals; see below for further analysis of other behavioral parameters). We first analyzed the effects of varying stimulus number. A single stimulus did not lead to speeding afterwards, but increasing the number of stimuli, even to 2 or 3 stimuli, led to a scalable increase in speed lasting over a minute afterwards (Fig. 1g). The magnitude of the speed increase scaled proportionately with the number of stimuli, saturating after ∼15 ASH stimuli (Fig. 1g). An analysis of individual animals revealed that this scalable increase was due to individual animals varying their locomotion speeds in a graded manner in response to different stimulus patterns (as opposed to animals exhibiting a binary speeding response that was more likely to occur with more stimuli; Extended Data Figure 1d-f). Provided that ASH was stimulated two or more times, the decay rate of the animals’ speed back to baseline was similar for different numbers of stimuli (Fig. 1g, right). This indicates that once the animals have received multiple ASH stimuli, they elevate their speed proportional the number of stimuli detected, and then speed gradually decays back to baseline over minutes.

We next examined how close successive ASH stimuli need to be spaced together in time in order to cause this long-lasting speed increase. To do so, we varied the interstimulus interval between optogenetic stimuli. Since single stimuli were ineffective at driving speeding, we reasoned that there should be some interstimulus interval beyond which speeding does not occur, i.e. where every stimulus is perceived as an isolated stimulus. We found that stimuli spaced apart by 32 sec were still integrated to increase animal speed (Fig. 1h). However, stimuli were not integrated when spaced >60 sec apart (Fig. 1h). Together, these data suggest that when multiple ASH stimuli are placed within half a minute of one another, it induces persistent speeding that then decays over minutes.

These results suggested that once an animal has received multiple ASH stimuli, it enters into a stable aversive behavioral state lasting minutes, during which time animals integrate stimuli to scale their speed. To further examine the temporal properties of this sensory input integration, we examined how far apart groups of ASH stimuli could be spaced and still be integrated to increase animal speed. For this, we delivered two groups of 5x ASH stimuli, and varied the time lag between the two groups (Fig. 1j). Even when the two groups of stimuli were spaced apart by 2.5 minutes, they could still be integrated: the response to the second group of stimuli was stronger due to prior ASH activation 2.5 minutes ago (Fig. 1j). Thus, multiple ASH stimuli induce entry into a behavioral state that persists for minutes. During this behavioral state, animals scale their speed based on the number of ASH stimuli that have been detected over minutes.

We next asked if a sensory evidence accumulation framework could explain the observed variation in speed dynamics across stimulus patterns. We built a computational model where instantaneous speed is predicted by the integrated sum of past stimuli weighted by an exponential decay function (Fig. 1k). We evaluated this model using mean square error of the model predictions, comparing it to multiple alternatives that differ in how they weight past stimuli: (1) integration with a linear (rather than exponential) decay; (2) uniform integration over a fixed time window in the past; (3) simple averaging of recent stimuli (i.e. gaussian smoothing); and (4) instantaneous responses to stimuli (i.e. a null model with no integration). Models were fit by minimizing error on a training dataset consisting of recordings with interstimulus intervals of 4, 16, and 64 seconds. The models were validated on 32-second interval data and tested on datasets with 8 sec interstimulus intervals, as well as a grouped stimulus dataset (Fig. 1k-m, Extended Data Figure 1g). All models with integration of past stimuli performed better than the null model with no integration, with the exponential decay model achieving the best fit for the data. The estimated decay constant of this model corresponded to a half-decay of 95 seconds (Extended Data Figure 1g). We systematically degraded the model parameters (the exponential decay rate λ and the stimulus coefficient c_s_ that scales response magnitude) and found that changing the decay parameter in any direction substantially degraded model performance (Fig. 1m), suggesting that the long decay rate was important for model performance. These results argue that the behavioral variation is well explained by a model where ASH stimuli are integrated over minutes to change behavior.

Having established the kinetics of ASH stimulus integration with this optogenetic paradigm, we next switched our focus to examining whether repeated encounters with a natural ASH stimulus could similarly evoke this aversive behavioral state. To test this, we place animals on a circular pad of NGM agar that was surrounded by NGM agar containing copper chloride, a chemical that activates ASH and triggers reversals (Extended Data Figure 1h-k). Bacterial food was uniform across the entire arena, including the area with copper. Animals explored these arenas and routinely ventured into the copper zone. When they approached the copper boundary, they reliably exhibited reversals (Extended Data Figure 1i). We measured how animal speed changes as a function of recent copper encounters. Indeed, the copper encounters led to a speeding effect afterwards, with a larger magnitude increase in animals that encountered copper many times in close succession (Extended Data Figure 1l-m). These results indicate that repeated stimulation with the natural ASH stimulus copper leads to increased speed, matching the optogenetics results.

Finally, we sought to examine whether the aversive behavioral state induced by repeated ASH stimulation impacted how animals respond to other sensory stimuli. For this, we utilized a dual optogenetics setup, where ASH was activated by the red-shifted opsin Chrimson, after which the appetitive sensory neuron AWA was activated using the blue-shifted opsin CoChR. We found that repeated ASH stimulation did not change either the magnitude or persistence of behavioral responses to AWA activation (Fig. 1q; controls in Extended Data Figure 1n). In contrast, activating ASH repeatedly led to more persistent responses to future ASH stimuli (Fig. 1p-q, Extended Data Figure 1n). This suggests that the ASH-induced state sensitizes animals to subsequent ASH stimuli, but not necessarily other sensory inputs. Taken together, this full set of results characterizes an aversive behavioral state that is triggered by repeated ASH stimulation. This state occurs when animals have detected multiple ASH stimuli and consists of multiple behavioral changes: increased speed, as well as reduced feeding and defecation. The properties of the state are scaled by the number of ASH stimuli detected over minutes, with more ASH stimuli leading to a more robust behavioral change.

### Sensory integration occurs downstream of ASH calcium, but is not a response to repeated reversals

We next began to examine where in the nervous system integration of ASH stimuli might be taking place. We first tested whether ASH itself displays persistent activity after it is repeatedly stimulated. We expressed the red light-activated opsin Chrimson under the ASH-specific promoter and delivered 2-second light pulses every 10 seconds to freely-moving animals on food, while simultaneously measuring ASH calcium dynamics with GCaMP (Fig. 1n). Repeated ASH photoactivation produced transient calcium responses that decayed back to baseline within 10 seconds (Fig. 1o). Repeated ASH photoactivation did not potentiate ASH calcium responses beyond the first stimulus (Fig. 1o). This indicates that ASH does not exhibit persistent neural activity and suggests that integration takes place somewhere downstream of ASH calcium influx. These results do not exclude signaling changes in ASH downstream of calcium influx, accumulation of ASH transmitters, or integration by downstream neurons.

Given that ASH induces a reversal each time it is stimulated, we asked whether directly stimulating the reversal circuit could induce the aversive behavioral state. For this, we repeatedly activated either AIB, AVA, or RIM, three well-established reversal-promoting interneurons^30,31,33,39^. While this reliably induced a reversal with each optogenetic stimulus, there was no elevation in speed during inter-stimulus intervals or after the last stimulus (Fig. 1i). Thus, repeatedly inducing reversals via activation of the reversal circuit cannot elicit the behavioral state. In addition, repeatedly activating RIB, a forward-promoting neuron^40^, caused immediate speeding during optogenetic stimulation, but no persistent behavioral change (Fig. 1i). This suggests that causing animals to speed up via a downstream network perturbation does not automatically induce a stable behavioral response. Taken together, these results suggest that integration occurs downstream of ASH calcium, but activation of the reversal circuit is not sufficient for integration. After excluding these possibilities, the most parsimonious remaining explanation was that other neurons downstream of ASH were performing this long timescale integration. Therefore, we next took advantage of population calcium imaging technologies available in *C. elegans* to perform a brain-wide activity screen for neural integrators downstream of ASH.

### Brain-wide neural recordings identify three interneurons – ADA, AVH, and PVQ – that act as neural integrators downstream of ASH

To identify candidate integrators of aversive stimuli, we performed whole-brain calcium imaging in freely-moving animals. Animals were recorded while exploring arenas with standard NGM agar surrounded by NGM containing copper (Fig. 2a; as described above; bacterial food was uniform across entire arena, including copper area). We imaged transgenic animals expressing pan-neuronal NLS-GCaMP7f and the NeuroPAL transgene, which provides fluorescent barcoding for neuron identification^14^ (Fig. 2a). GCaMP was normalized to the pan-neural TagRFP in NeuroPAL. Animals were recorded for 16 minutes using our previously described live-tracking microscope^19,41^. At the end of each recording, they were immobilized and multi-spectral images were collected to identify neuron classes with NeuroPAL. All image processing for trace extraction and neuron identification was performed using automated software, as previously described^41,42^. Using this approach, we recorded a total of 27 animals with 96 neurons recorded on average per animal.

**Figure 2.**
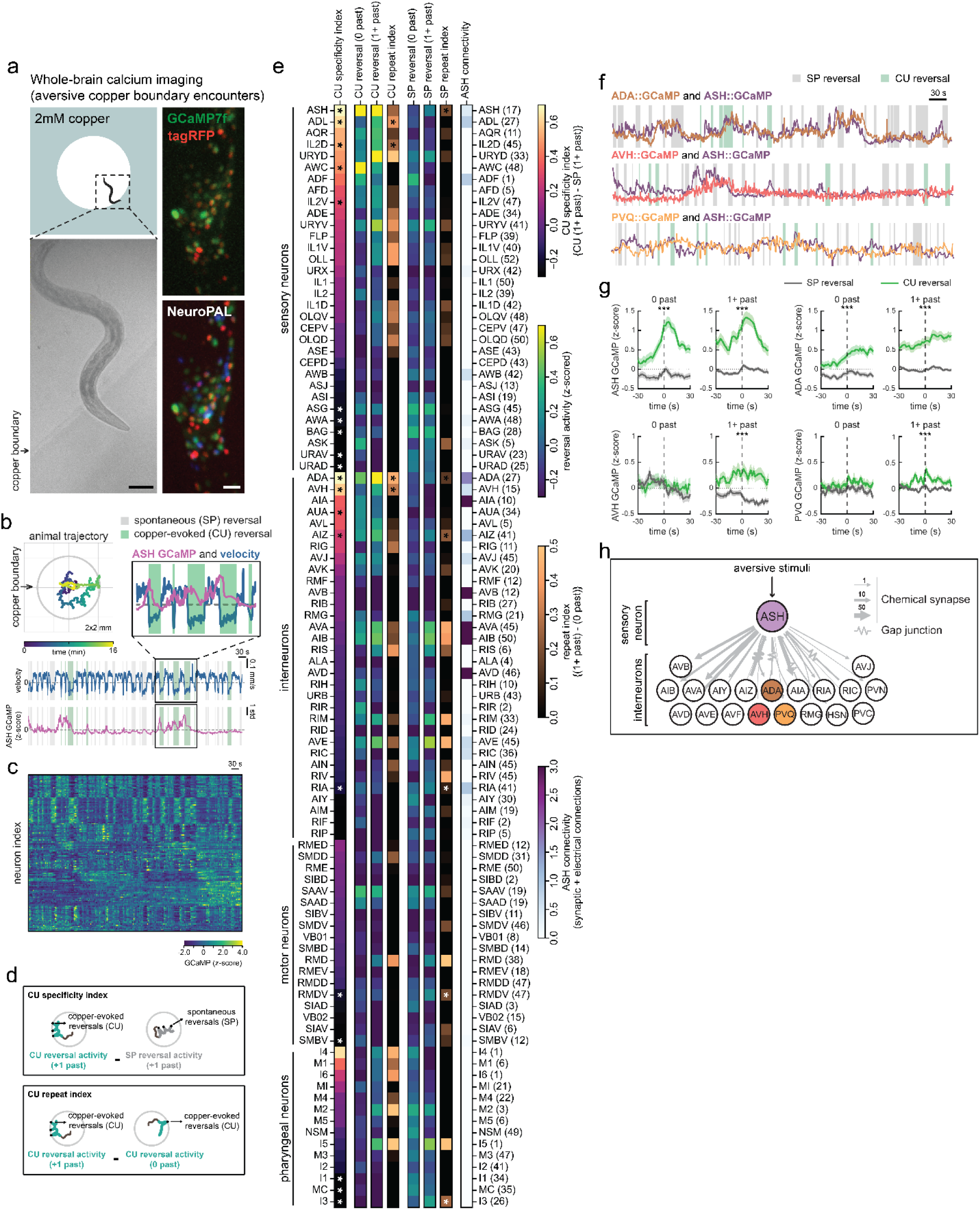
Brain-wide calcium imaging in freely-moving *C. elegans* identifies integrator neurons that stably accumulate information about recent aversive sensory encounters. a. Schematic of brain-wide neural calcium recordings during repeated encounters with a naturally aversive copper-infused agar boundary, and representative images of GCaMP/TagRFP and NeuroPAL from a single animal. In the upper right image, the copper area is false-colored blue for illustrative purposes. b. Movement trajectory of a representative animal during repeated encounters with the boundary, showing velocity (top) and z-scored ASH activity (bottom). c. All recorded GCaMP traces across the brain from the representative animal, shown over the same time stretch. d. Two metrics to help identify integrator neurons: (1) copper (CU) specificity index, which denotes relative calcium dynamics during repeated copper-evoked (CU) vs repeated spontaneous (SP) reversals; and (2) copper (CU) repeat index, which denotes the relative calcium dynamics after repeated copper-evoked reversals vs after an isolated CU-evoked reversal. e. Average neuronal activity during copper-evoked (CU) and spontaneous (SP) reversals, categorized by the number of preceding reversals of the same type within a 30 second time window. Neurons were ranked according to their copper specificity index, which indicates differences between repeated CU vs SP reversals (*p<0.05, Bonferroni-corrected two-tailed Mann-Whitney test). Neurons with high copper specificity were also tested for CU and SP repeat indices, which indicates differences between repeated vs single reversals of the same type (*p<0.05, one-tailed Mann-Whitney test). Also shown is the average number of synaptic and electrical ASH inputs received by each neuron. The numbers in parentheses following neuron names indicate the number of recordings made for that neuron across all datasets. f. Representative changes in the activity of ADA, AVH and PVQ neurons during repeated encounters with the copper boundary. The top two traces are from whole-brain recordings, whereas the bottom trace (PVQ+ASH) is from a sra-6::GCaMP recording in the same arenas used for whole-brain calcium imaging. g. Event-triggered averages of ADA, AVH and PVQ neurons during copper-evoked (CU) and spontaneous (SP) reversals, categorized by the number of preceding reversals of the same type within a 30 second window. For example, “0 past” indicates no previous reversals of that type in the last 30 seconds, whereas “1+ past” indicates one or more reversals (of that type) within the last 30 seconds. ***p<0.001, one-tailed Mann-Whitney test. h. A circuit diagram showing all neurons that lie downstream of ASH in *C. elegans* connectome.

To characterize these datasets (example in Fig. 2b-c), we first examined the behavior of the recorded animals. Encounters with the copper boundary elicited reversals (Fig. 2a and Extended Data Figure 2a) and repeated encounters caused animals to elevate their speed (Extended Data Figure 2b). We also asked which sensory neurons showed altered activity upon copper encounter. We ranked neurons by a “copper specificity index”, which quantifies the increase in activity during copper-induced reversals versus spontaneous reversals (Fig. 2d). Among sensory neurons, ASH had the highest copper specificity, followed by weaker copper responses in ADL, IL2D, AWC, and IL2V (Fig. 2e). Thus, in these whole-brain recordings, encounters with copper activate ASH, trigger reversals, and induce stable speeding.

We next analyzed all of the recorded neurons in an effort to identify specific neurons with neural integrator properties. We reasoned that integrator neurons should fulfill two criteria. First, they should show higher activity during copper-evoked reversals versus spontaneous reversals. We used the “copper specificity index” described above that quantifies this (Fig. 2d). Second, integrator neurons should show higher activity when there are repeated copper encounters in close succession (within 30 seconds of each other). We devised a “copper repeat index” that quantifies this (Fig. 2d). We examined all neurons recorded across all animals and identified four neuron types where both indices had significant positive values (Fig. 2e shows all neurons; see figure legend for multiple comparison correction statistics). Two of these neurons were sensory neurons (ADL and IL2D) that had relatively weak responses to copper. The other two were interneurons, ADA and AVH, that had strong responses to repeated copper encounters.

Two other interneurons (AUA and AIZ) showed responses to copper encounters, but did not exhibit accumulating activity during repeated copper encounters. These results raised the possibility that ADA and AVH may act as neural integrators.

The above analyses identified candidate integrator neurons based on their responses to copper encounters. Given that ASH and the candidate neural integrators were co-recorded in animals, we also directly examined whether ADA and AVH dynamics indeed tracked ASH activity in these datasets. Indeed, both neurons showed neural activity closely aligned to ASH, though their activities showed smoother signals that generally lagged behind ASH (Fig. 2f-g).

ADA and AVH both receive direct synaptic inputs from ASH; ADA is also connected to ASH via gap junctions. Since our brain-wide imaging approach misses neurons in the tail, we also considered ASH synaptic targets whose cell bodies are in the tail. Among these neurons (PVC, PVN, and PVQ), PVQ is unique in that it receives ASH synaptic inputs and is also connected to ADA via gap junctions. Therefore, we performed targeted calcium imaging of ASH and PVQ neurons during copper boundary encounters. PVQ neurons also had significant copper encounter responses that scaled with repeated encounters (Fig. 2f-g). Together, these results identify ADA, AVH, and PVQ as candidate interneurons downstream of ASH that integrate aversive experiences (Fig. 2h).

We next directly examined if these neurons could integrate ASH activity and display persistent activity. We expressed Chrimson in ASH neurons and delivered 2-second light pulses at 10-second intervals while simultaneously recording calcium dynamics in ADA, AVH, or PVQ neurons in freely-moving animals using cytoplasmic GCaMP7f (Fig. 3a-c). Repeated ASH photoactivation produced elevated calcium responses above baseline in all tested neurons (Fig. 3a-c). AVH neurons showed significantly higher activity following repeated ASH stimuli, with persistent responses lasting >60 seconds after stimulus offset. PVQ neurons similarly showed elevated activity following repeated stimulation, with persistent responses lasting ∼60 seconds after stimulus offset. ADA responses were also detected, though they were more transient, decaying after ∼10 seconds. These results suggest that AVH and PVQ neurons integrate ASH inputs through ramping activity and persistent responses; ADA integrates ASH inputs on a faster timescale. These responses appear to be fairly unique among neurons in the *C. elegans* nervous system, based on our global analysis of whole-brain activity showing that other recorded neurons did not exhibit these stable responses to aversive sensory encounters.

**Figure 3.**
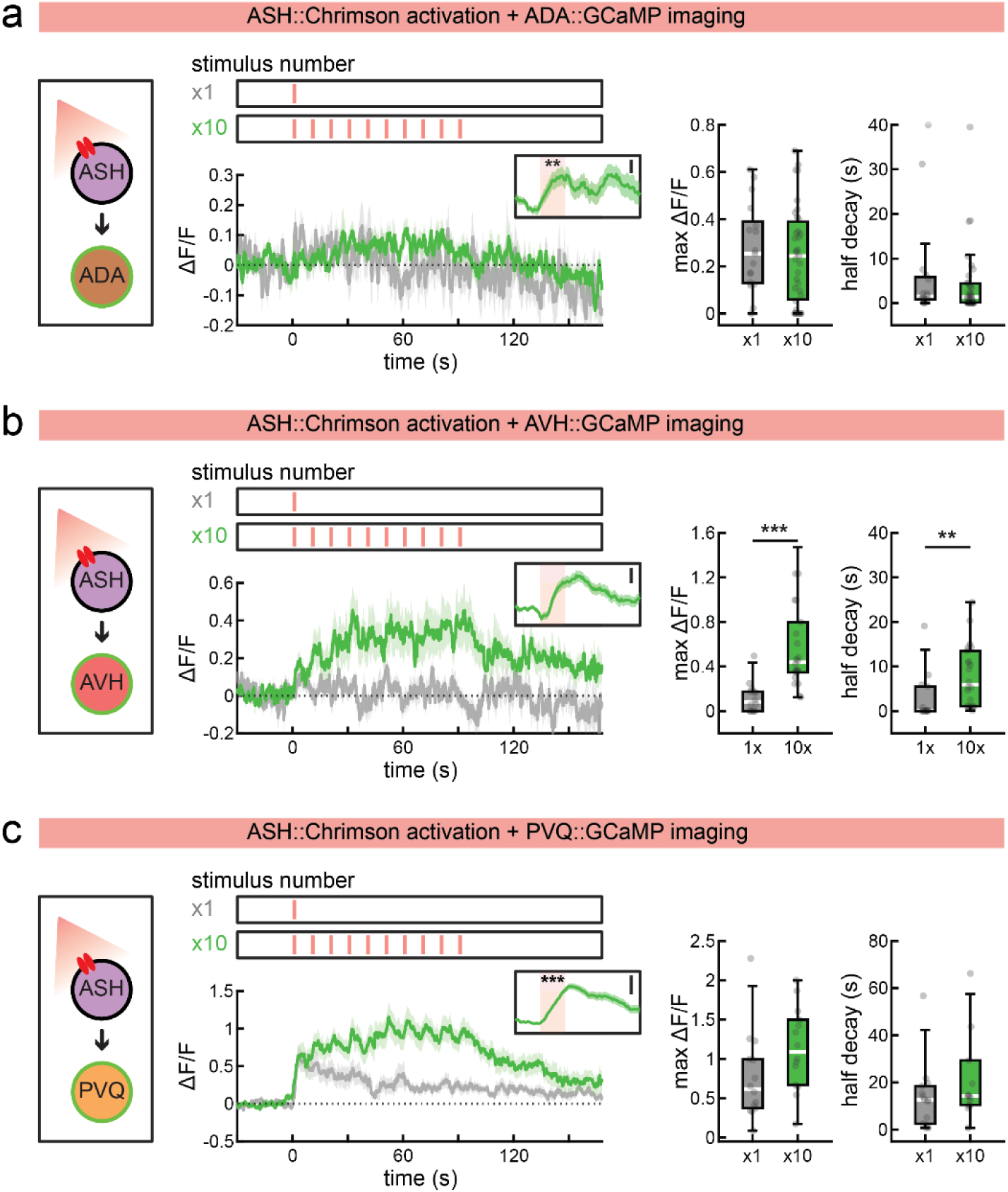
Optogenetic activation of ASH elicits persistent calcium responses in the putative integrator neurons ADA, AVH, and PVQ. a. Calcium responses in ADA neurons recorded during optogenetic activation of ASH. Left: event-triggered averages of normalized ASH calcium dynamics after a single or repeated ASH stimulation. The inset shows averages from −2s to 8s across all 10 stimuli, normalized to the pre-stimulus baseline (−2s to 0s; **p<0.01, Wilcoxon signed-ranked test); Middle: maximum change in calcium levels at last stimulus; Right: half decay time of calcium levels after last stimulus offset. Boxes show the median and interquartile range (25th-75th percentile). Whiskers extend to the most extreme value within 1.5x the interquartile range. Dots are individual animals. Inset scale bars: 0.01 a.u. n=16 animals for x1; n=35 animals for x10. b. Calcium responses in AVH neurons recorded during optogenetic activation of ASH, shown as in (a). Middle: maximum change in calcium levels at last stimulus offset (**p<0.01, Wilcoxon rank-sum test); Right: half decay time of calcium levels after last stimulus offset (*p<0.05, Wilcoxon rank-sum test). Inset scale bars: 0.03 a.u. n=14 animals for x1; n=22 animals for x10. c. Calcium responses in PVQ neurons recorded during optogenetic activation of ASH, shown as in (a) (***p<0.001, Wilcoxon signed-ranked test). Inset scale bars: 0.1 a.u. n=14 animals for x1 and x10.

### The integrator neurons ADA, AVH, and PVQ functionally contribute to the aversive behavioral state, controlling distinct behavioral features

Our calcium imaging results suggest that ADA, AVH, and PVQ exhibit dynamics typical of neural integrators, both during natural sensory stimulus encounters and during optogenetic ASH activation. We next wanted to test whether these neurons functionally contribute to the ASH-induced aversive behavioral state. AVH has been implicated in behavioral responses to touch^43^, but otherwise these three interneurons do not have any established behavioral functions in *C. elegans*. To address this, we selected cell-specific promoters for each neuron (Extended Data Figure 2c) so that we could perturb their activity with transgenic tools and examine behavior. We repeatedly photoactivated ASH neurons while hyperpolarizing ADA, AVH or PVQ neurons with a leaky potassium channel *unc-103(gf)* (Fig. 4a). All manipulations impaired the ASH-induced speeding response relative to wild type controls (Fig. 4b), though to different degrees (baseline speed was not diminished by silencing any of these neurons, Extended Data Figure 3a). Silencing ADA or AVH dramatically impaired the ASH-induced speeding response, whereas silencing PVQ effectively increased the number of ASH stimuli that were required for animals to exhibit elevated speed. The immediate ASH-induced reversal response was unaffected by silencing any of these three neurons (Fig. 4c; Extended Data Figure 3b-d). Further, hyperpolarization of each interneuron impaired the reductions in feeding and defecation rates caused by repeated ASH stimulation (Fig. 4c; Extended Data Figure 3b-d). These results suggest that each of these three neurons contributes to the ASH-induced behavioral state, with ADA and AVH playing a key role in speeding and PVQ controlling the rate of onset of the ASH-induced aversive state.

**Figure 4.**
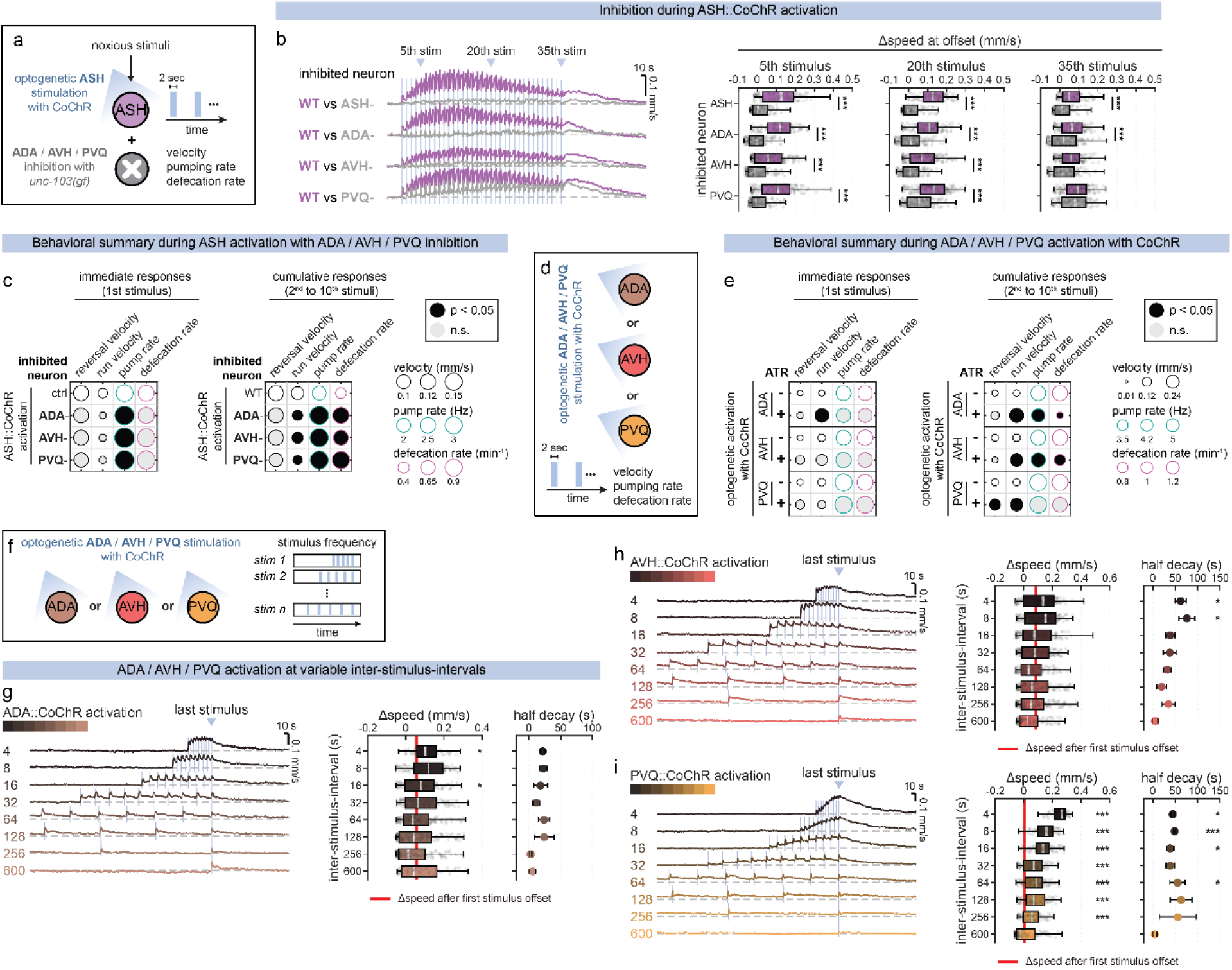
The identified integrator neurons control different aspects of behavior during the aversive behavioral state. a. Schematic of optogenetic ASH neuron activation in transgenic animals with inactivated ASH, ADA, AVH or PVQ neurons, and simultaneous behavioral recordings in freely-moving animals. Noxious stimuli (harsh touch, heavy metals, aversive odors) endogenously activate ASH neurons. To mimic this activation, animals expressing CoChR in ASH received repeated 2-second light pulses; velocity, pumping, and defecation frequency were assessed. ASH was inactivated via tetanus toxin expression, whereas ADA, AVH, and PVQ were inactivated via expression of the leaky potassium channel UNC-103(gf). b. Left: Event-triggered changes in locomotion speed after repeated optogenetic stimulation of ASH neurons in transgenic animals with inactivated ASH, ADA, AVH or PVQ neurons. Right: Changes in speed after 5^th^, 20^th^, and 35^th^ stimulus offset. Each behavior compared to day-matched wild-type controls (purple). ***p<0.001, Bonferroni-corrected empirical p-value for speed changes at stimulation offset. n = 2-4 recorded plates with 27-64 animals per plate. c. Summary of behavioral changes during optogenetic ASH neuron activation in transgenic animals with inactivated ADA, AVH or PVQ neurons. Mean reversal and run velocity were measured from the stimulus and post-stimulus periods, respectively; mean pumping and defecation rates were measured from the combined period. Each behavior compared to day-matched wild-type controls. *p<0.05, Bonferroni-corrected Wilcoxon rank-sum test. n=15-17 animals for each genotype. d. Schematic of optogenetic activation of ADA, AVH or PVQ neurons with CoChR. e. Summary of behavioral changes during optogenetic activation of ADA, AVH or PVQ neurons. Each behavior compared to day-matched wild-type controls. *p<0.05, Bonferroni-corrected Wilcoxon rank-sum test. n=12-36 animals for each genotype. f. Schematic of different optogenetic neuron activation patterns for ADA, AVH or PVQ neurons. Variable-frequency: 2-second stimuli delivered with 4, 8, 16, 32, 64, 128, 256, or 600 second inter-stimulus-intervals for a total of 10 times. g. Event-triggered changes in speed after repeated optogenetic stimulation of ADA at different inter-stimulus-intervals. Stimulus design and analysis the same as in Fig. 1h. Boxplot shows the average change in speed at the last stimulus offset (*p<0.05, Bonferroni-corrected empirical p-value for speed at stimulation offset compared to the speed after the first stimulus). Dot plot shows the half decay time of the average speed after the last stimulus offset (*p<0.05, Bonferroni-corrected empirical p-value for change in half decay times compared to the longest inter-stimulus-interval condition (600 seconds)). n = 1-3 recorded plates with 30-75 animals per plate. h. Event-triggered changes in speed after repeated optogenetic stimulation of AVH, shown as in (g). *p<0.05, Bonferroni-corrected empirical p-value. n = 3 recorded plates with 31-79 animals per plate. i. Event-triggered changes in speed after repeated optogenetic stimulation of PVQ, shown as in (g). *p<0.05, ***p<0.01, Bonferroni-corrected empirical p-value. n = 3-4 recorded plates with 22-36 animals per plate.

We next tested whether activation of these integrator neurons is sufficient to drive long-term behavioral changes. We expressed CoChR in each neuron and delivered repeated 2-second stimuli at varying interstimulus intervals (Fig. 4d). This allowed us to examine behavioral responses to a single optogenetic stimulus (the first stimulus), as well as repeated stimuli spaced apart by different time intervals. Stimulation of ADA or AVH led to robust and immediate speeding, elevating the animal’s speed by ∼0.1mm/sec (Fig. 4e; Extended Data Figure 3e-f). The speeding observed in response to a single stimulus was only mildly increased when several stimuli were delivered in close succession (Fig. 4f-h). In addition, activation of ADA or AVH reduced feeding and defecation rates, which are both behavioral hallmarks of the ASH-induced state. After stimulation, behavior decayed back to baseline with a half-decay time of ∼30-60 sec. Thus, ADA or AVH activation induces immediate and long-lasting speeding and other behavioral changes.

The results were quite different for optogenetic activation of the other integrator neuron, PVQ. A single optogenetic stimulation of PVQ did not elevate the animal’s speed, feeding, or defecation (Fig. 4e; Extended Data Figure 3g). However, repeatedly activating PVQ did eventually elevate the animals’ speed after 3-4 successive stimuli. The elevated speed decayed very slowly, lasting minutes. Consistent with this, single PVQ stimuli were integrated even when spaced apart by >4 min (Fig. 4i). These results indicate that PVQ stimulation does not cause an immediate behavioral state change. Instead it gradually increases the animal’s speed when it is repeatedly activated within a minutes-long time window.

Overall, these results suggest that ADA and AVH can directly drive the speeding observed in the aversive behavioral state. In contrast, PVQ does not directly drive immediate speeding. Instead, it can only drive speeding if it has been recently activated several times. Together with the neural silencing results above, this raised the possibility that, in contrast to ADA and AVH, PVQ activation might facilitate stronger responses to subsequent stimuli, perhaps helping induce entry into the persistent behavioral state, which then consists of persistent speeding scaled by the number of ASH stimuli. To directly test this, we examined whether stimulation of PVQ could elicit a priming effect, where it could enhance behavioral responses to subsequent ASH activation (Fig. 5a-d). For this, we used dual optogenetics, stimulating PVQ five times (using blue light-activated CoChR) and then subsequently stimulating ASH five times (using red light-activated Chrimson) (Fig. 5d). We tested multiple time lags between stimulation of the two neurons. As a point of comparison, we performed the same dual optogenetics experiments on ADA and AVH (Fig. 5b-c). Speed dynamics were compared to control animals expressing ASH::Chrimson alone, which received identical light stimulation (controls for dual optogenetic experiments in Extended Data Figure 4a-c). Activation of ADA or AVH led to immediate speeding. However, the behavioral responses to subsequent ASH stimulation were not enhanced, when the speed was still decaying back to baseline or afterwards when speed had fully decayed (Fig. 5b-c). Thus, ADA and AVH activation induces speeding, but does not enhance responses to subsequent ASH stimuli. By contrast, PVQ activation notably enhanced subsequent ASH-evoked speed responses, even at delays >4 minutes later when speed had decayed back to baseline (Fig. 5d; the strains being compared had the same responses to ASH stimulation with red light when there was no prior blue light stimulation, Extended Data Figure 4b). Taken together, these results suggest that ADA and AVH integrate ASH activity and drive immediate speeding, proportional to the number of ASH stimuli detected. In contrast, PVQ integrates ASH activity to set a long-lasting internal state during which animals display stronger responses to incoming ASH stimuli.

**Figure 5.**
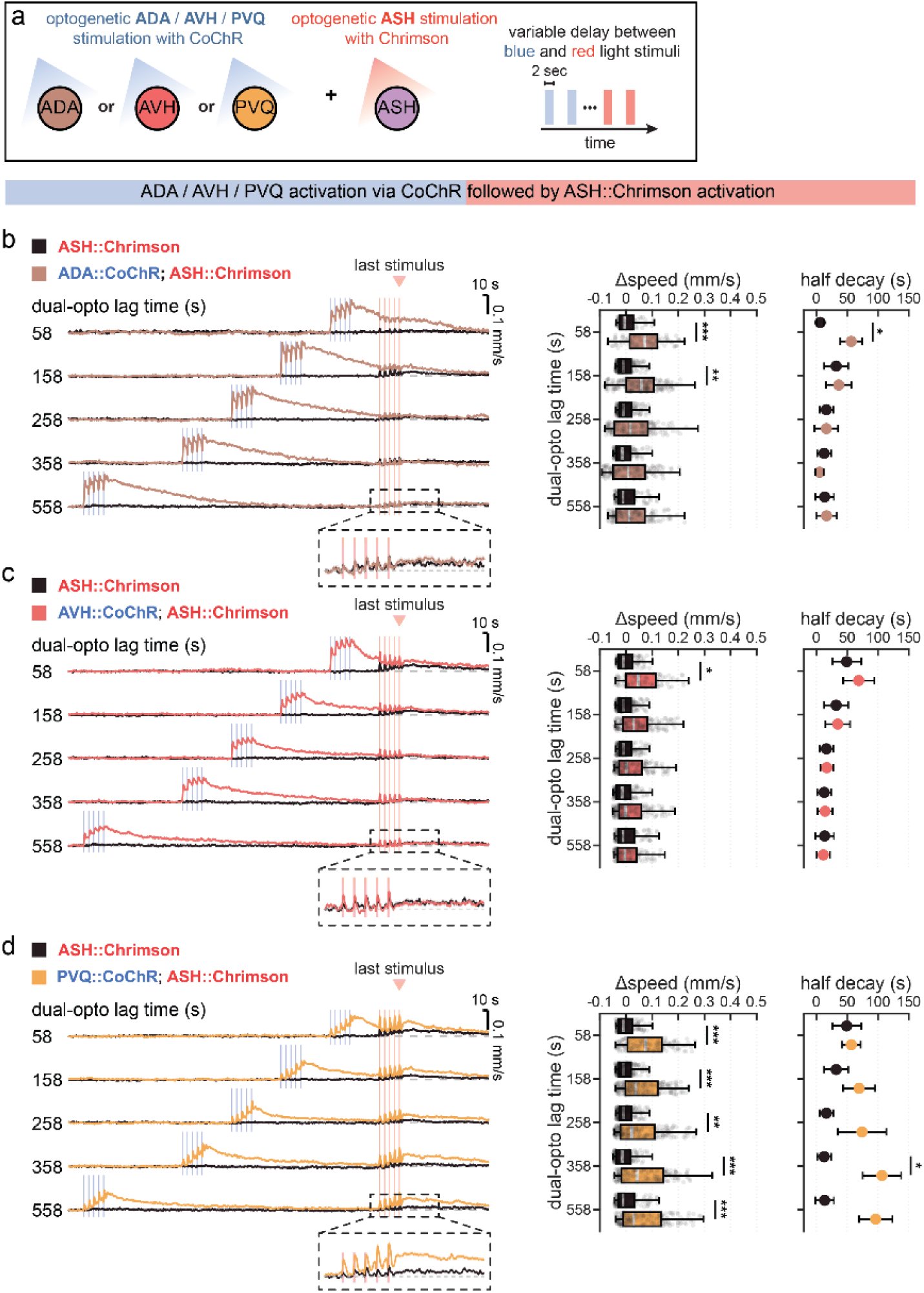
The identified integrator neurons differentially regulate responsiveness to future aversive stimuli. a. Schematic of dual-color optogenetic activation of ADA, AVH or PVQ neurons with CoChR, followed by optogenetic activation of ASH with Chrimson. b. Left: Event-trigger changes in speed after dual-color optogenetic activation of ADA with blue light, followed by delayed optogenetic activation of ASH neurons with red light. Right: Box plot shows the average change in speed at the last red stimulus offset (**p<0.01, ***p<0.001, Bonferroni-corrected empirical p-value compared to ASH::Chrimson animals that received identical light). Dot plot shows the half decay time of the average speed after the last stimulus offset (*p<0.05, Bonferroni-corrected empirical p-value for change in half decay times compared to ASH::Chrimson animals that received identical light). n = 2-4 recorded plates with 19-117 animals per plate. c. Dual optogenetic results for activation of AVH, followed by ASH, shown as in (k). *p<0.05, Bonferroni-corrected empirical p-value. n = 2-4 recorded plates with 22-103 animals per plate. d. Dual optogenetic results for activation of PVQ, followed by ASH, shown as in (k). *p<0.05, **p<0.01, ***p<0.001, Bonferroni-corrected empirical p-value .n = 2-4 recorded plates with 22-103 animals per plate.

### The integrator neurons AVH and PVQ act in parallel, using different integration mechanisms and different transmitters

We next examined the mechanisms by which these neural integrators function within the *C. elegans* nervous system. At one extreme, they could all function together and comprise a single integrator circuit. At the other extreme, they could all operate independently. The results above suggested independent functions, at least at the level of behavioral control. To more directly address this, we first examined whether the integrator neurons depend on one another for their function. We repeatedly activated one integrator neuron with CoChR while silencing another with *unc-103(gf)*, and performed reciprocal experiments (Fig. 6a). We then quantified the speed increase for each of these conditions. AVH and PVQ neurons were required for the speed responses to ADA activation, suggesting AVH and PVQ neurons act downstream of ADA (Fig. 6b-c). However, activation of AVH or PVQ led to behavioral changes that did not depend on the other integrators (Fig. 6b-c; note that no genetic silencing lines impaired the effects of AVH or PVQ stimulation). This suggests that AVH and PVQ are functionally downstream of ADA and can function in parallel to one another.

**Figure 6.**
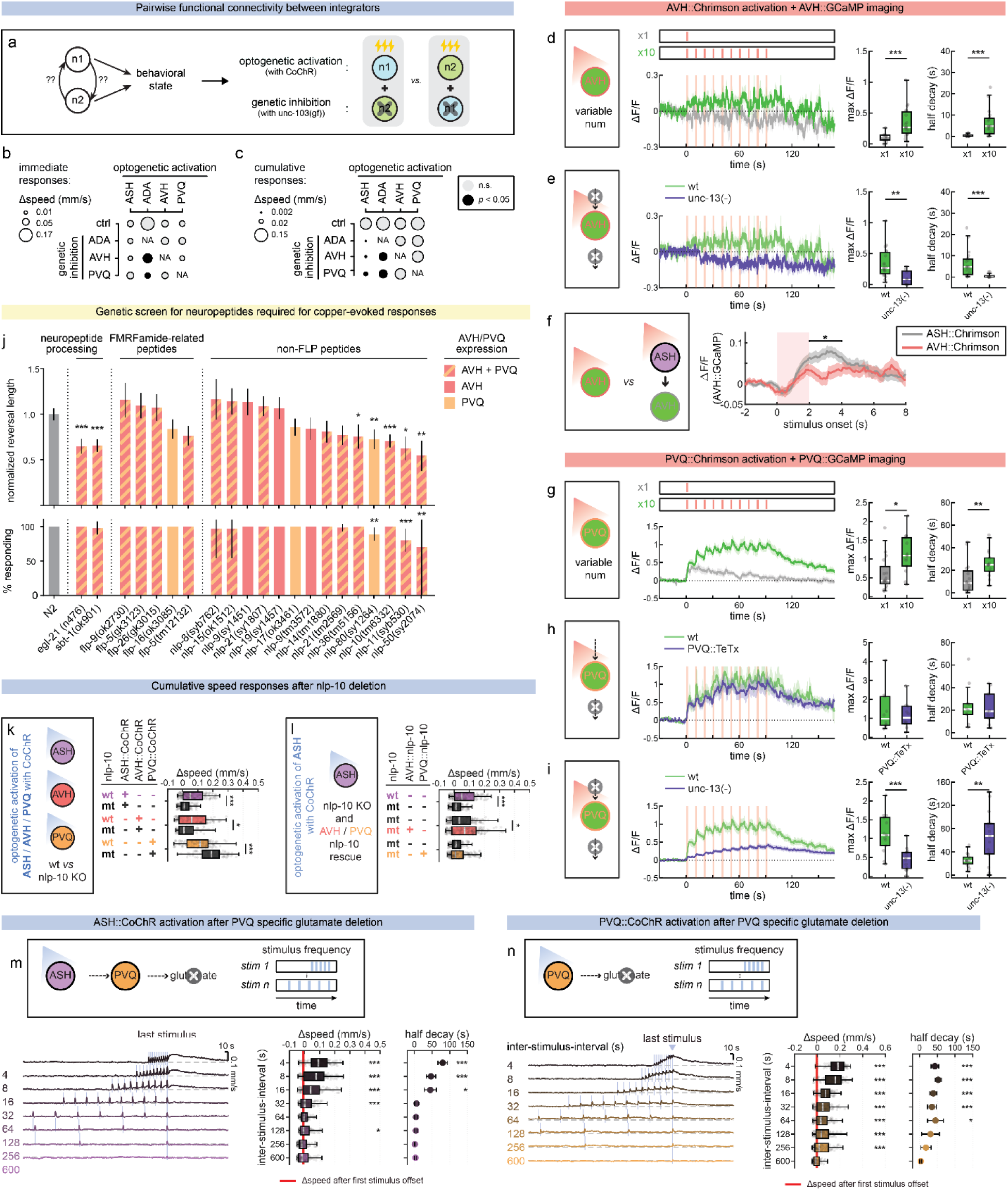
The integrator neurons function in parallel to one another and employ different mechanisms for persistent activity and controlling behavior. a. Schematic of the genetic approaches used to test pairwise functional relationships between ASH, ADA, AVH and PVQ neurons. One neuron in the group is optogenetically stimulated while another is silenced, and the effects on behavior is measured. All combinations are tested. b. Summary of pairwise relationships between ASH, ADA, AVH and PVQ neurons. This plot shows immediate changes in speed (i.e. after first optogenetic stimulus). Circle size indicates average speed and color indicates Bonferroni-corrected empirical p-value. For neuronal inhibitions during ASH::CoChR activation, n = 3-4 recorded plates with 27-64 animals per plate. For neuronal inhibitions during ADA::CoChR activation, n = 5 recorded plates with 29-78 animals per plate. For neuronal inhibitions during AVH::CoChR activation, n = 3-4 recorded plates with 28-56 animals per plate. For neuronal inhibitions during PVQ::CoChR activation, n = 4 recorded plates with 26-56 animals per plate. c. Summary of pairwise relationships between ASH, ADA, AVH and PVQ neurons. This plot shows cumulative changes in speed (i.e. after the 5th optogenetic stimulus). Circle size indicates average speed and color indicates Bonferroni-corrected empirical p-value. d. Optogenetic AVH neuron activation and simultaneous AVH calcium recordings. Left: event-triggered averages of normalized AVH calcium dynamics after a single or repeated AVH::Chrimson stimulation; Middle: maximum change in calcium levels at last stimulus offset; Right: half decay time of GCaMP signal after last stimulus offset. Boxes show the median and interquartile range (25th-75th percentile). Whiskers extend to the most extreme value within 1.5x the interquartile range. Dots are individual animals. ***p<0.001, Mann-Whitney test. For x1, n=11 animals; for x10 n=19 animals. e. Optogenetic AVH neuron activation and simultaneous AVH calcium recordings in unc-13 synaptic mutants, displayed as in (d). **p<0.01, ***p<0.001, Mann-Whitney test. For *unc-13(-)*, n=9 animals; for wild-type controls (*wt*), n= 19 animals. f. Event-triggered changes in AVH neuron calcium activity during optogenetic activation of ASH or AVH neurons. Baseline-corrected to the 2-second pre-stimulus period. Same data as in (2h) and (4d). *p<0.05, Mann-Whitney test. g. Optogenetic PVQ neuron activation and simultaneous PVQ calcium recordings, displayed as in (d). *p<0.05, **p<0.01, Mann-Whitney test. For x1, n=16 animals; for x10 n=14 animals. h. Optogenetic PVQ neuron activation and simultaneous PVQ calcium recordings in animals expressing PVQ::TeTx, displayed as in (d). For *PVQ::TeTx*, n=10 animals; for wild-type controls (*wt*), n=10 animals. i. Optogenetic PVQ neuron activation and simultaneous PVQ calcium recordings in unc-13 synaptic mutants, displayed as in (d). **p<0.01, ***p<0.001, Mann-Whitney test. For *unc-13(-)*, n=15 animals; for wild-type controls (*wt*), n=14 animals. j. Analysis of neuropeptide signaling mutants in a copper response assay. The indicated mutants were tested in a copper drop assay. Plots show average reversal length in response to application of a copper drop (top) and percent responding animals (bottom). Mean with 95% CI. Reversal length was normalized to day-matched wild type controls. Neuropeptide genes were selected based on their expression in either AVH, PVQ, or both neurons. *p<0.05, **p<0.01, ***p<0.001, Kruskal-Wallis test with Dunn’s multiple correction. n = 2-4 trials, 11-16 animals per trial. k. Changes in speed after optogenetic activation of ASH/AVH/PVQ neurons in wild-type versus *nlp-10* mutant animals. ***p<0.001, empirical p-value Shown are the responses after delivery of 20x 2s stimuli at 10Hz. n = 1-4 recorded plates with 17-78 animals per plate. l. Changes in speed after optogenetic ASH neuron activation in *nlp-10* mutant animals expressing *nlp-10* cDNA either in AVH or in PVQ neurons. *p<0.05, ***p<0.001, empirical p-value. Shown are the responses after delivery of 20x 2s stimuli at 10Hz. n = 3-5 recorded plates with 24-72 animals per plate. m. Change in speed after ASH optogenetic activation at variable inter-stimulus-intervals in animals with PVQ-specific *eat-4* knockout. Compare to figure 1h, displayed in the same manner, and figure Extended Data Figure 5c. *p<0.05, ***p<0.001, empirical p-value. n = 3 recorded plates with 23-46 animals per plate. n. Change in speed after PVQ optogenetic activation at variable inter-stimulus-intervals in animals with PVQ-specific eat-4 knockout. Compare to figure 4i, displayed in the same manner, and figure Extended Data Figure 5d. *p<0.05, ***p<0.001, empirical p-value. N = 3 recorded plates with 25-49 animals per plate.

We next examined the neural mechanisms that underlie neural integrator function. Our results so far suggest that ADA performs fast timescale integration, is required for the aversive behavioral state, and is functionally upstream of AVH and PVQ, serving a gating function. In contrast, AVH and PVQ perform long timescale integration and act in parallel to control different aspects of the behavioral state. We focused our attention on the mechanisms underlying longer timescale integration of AVH and PVQ. Integrator neurons may sustain stable activity via cell-intrinsic mechanisms, circuit-level mechanisms, or both. To distinguish between these possibilities, we first directly activated each neuron while recording it. Repeated AVH photoactivation with Chrimson led to an increase in AVH GCaMP (Fig. 6d). However, AVH activity was more transient after AVH stimulation compared to when we used ASH stimulation to activate AVH (Fig. 6f; and compare to Fig. 3b). In addition, AVH calcium responses were impaired in an *unc-13* mutant background with a global impairment in synaptic activity (Fig. 6e). These results suggest that AVH does not have cell-intrinsic persistent activity, but instead relies on other circuit elements triggered by ASH. In contrast, repeated PVQ photoactivation led to cumulative and persistent increases in PVQ GCaMP, matching the results from ASH-driven PVQ activation (Fig. 6g). To test whether this effect required PVQ outputs, for example due to a recurrently connected circuit, we repeated this experiment while impairing PVQ transmission via PVQ tetanus toxin expression. This had no impact on PVQ activation, suggesting this its output is not required to maintain its persistent activity (Fig. 6h). Moreover, PVQ even displayed persistent activity in an *unc-13* mutant background with a global impairment in synaptic activity across the entire nervous system (Fig. 6i; though we note that the amplitude of its activation was reduced suggesting that circuit activity contributes to the magnitude of activation). Taken together, these results suggest that persistent activity in AVH requires activation of other circuit elements during ASH stimulation, whereas PVQ is able to rely on cell-intrinsic mechanisms for its persistent activity.

AVH and PVQ neurons are enriched in dense core vesicles^7,44^, suggesting that they may use neuropeptidergic outputs to control these long timescale behaviors. These neurons express numerous neuropeptides and neuropeptide-processing enzymes^44^. AVH does not produce any classical neurotransmitters^45^, while PVQ has been reported to be glutamatergic^46^. To identify specific neuropeptides involved, we obtained mutants lacking AVH- and/or PVQ-expressed neuropeptides and screened them for a behavioral phenotype consistent with a deficit in speeding: intact copper-induced reversals, but slower movement during those reversals. Of these, *nlp-10* mutants showed the clearest phenotype of that type (Fig. 6j). To determine if NLP-10 is required for the aversive behavioral state, we measured speed dynamics in *nlp-10* mutants after ASH, AVH, or PVQ photoactivation. Loss of *nlp-10* impaired the cumulative speed increase after repeated ASH and AVH stimulation, but not PVQ stimulation (Fig. 6k). Loss of *nlp-10* did not impair baseline speed (Extended Data Figure 5a), ruling out a general locomotion deficit. To determine if *nlp-10* acts in AVH or PVQ neurons, we expressed *nlp-10* specifically in each neuron and tested cumulative speed responses after repeated ASH stimulation. Rescue of *nlp-10* specifically in AVH, but not PVQ, partially restored the aversion-induced speed increase (Fig. 6l). This suggests that AVH acts through the NLP-10 neuropeptide to induce speeding.

Finally, we also examined the transmitter output of PVQ that mediates its effects. We did not identify a PVQ-expressed neuropeptide with a behavioral phenotype and the only classical neurotransmitter reported to be used by PVQ is glutamate^46^. So, we examined whether cell-specific disruption of glutamate production by PVQ, via cell-specific deletion of the vesicular glutamate transporter *eat-4*^47^, impaired PVQ’s ability to induce the persistent behavioral state. Indeed, both the ASH- and PVQ-induced speeding effects were attenuated by PVQ glutamate deletion (Fig. 6m-n, Extended Data Figure 5b-e; the endogenous fluorescent reporter for *eat-4* expressed in PVQ, Extended Data Figure 5b, which was notable since reports of PVQ *eat-4* expression have been mixed^45,46^). These effects were most pronounced when stimuli were spaced far apart, suggesting that long timescale integration of successive PVQ stimuli is dependent on its glutamatergic output. Taken together, these results indicate that there are multiple integrator neurons in the *C. elegans* nervous system. ADA operates on the order of seconds and is required upstream of AVH and PVQ, serving a gating function. AVH and PVQ act over minutes, stably integrating ASH activation. Moreover, they utilize distinct integration mechanisms and act through distinct outputs to control distinct features of the aversive behavioral state.

## Discussion

Animals stably integrate information about their sensory surroundings to generate adaptive behavioral responses. Here, we identified and characterized neurons across the *C. elegans* brain that stably integrate aversive sensory experiences over minutes. We show that the dynamical properties of a set of newly identified integrator neurons – ADA, AVH, and PVQ – are critical for an aversive behavioral state. ADA is a fast timescale integrator that serves a gating function upstream of AVH and PVQ. AVH and PVQ exhibit minutes-long integration and act in parallel to control different aspects of the aversive state, such as high-speed locomotion versus priming the animal to increase responsiveness to future aversive stimuli. Our work suggests a model where aversive sensory information is processed in parallel streams by multiple integrators that ultimately combine to generate the full set of behavioral changes that comprise a global state.

The prevailing model for the control of behavioral states involves few-to-many signaling where a single neural hub integrates information and then induces a behavioral state through diffuse broadcasting to other neuron populations^48^. This architecture has been observed for mice, flies, and worms^49–53^. Our work here suggests that a different mechanism controls an integrative behavioral state that worms express in response to repeated aversive stimuli. Through brain-wide imaging, we identified multiple integrator neurons that are stably activated during repeated aversive stimulation. We then used cell-specific perturbations to show that the two long timescale integrators, AVH and PVQ, operate in parallel: their behavioral effects were unimpaired when the other was silenced. Moreover, they controlled different features of behavior, with AVH inducing stable speeding and PVQ inducing increased sensitivity to future aversive stimuli. This suggests that this aversive state can be decomposed to a set of behavioral features that map onto different neural substrates. This organization may allow for modular control over these distinct behavioral parameters to enable the flexible generation of a range of different behaviors. Previous work has shown how the neuropeptide tachykinin 2 can act independently in multiple regions of the mouse brain to similarly achieve modular behavioral control during social isolation stress^54^.

The aversive behavioral state we characterized here consists of multiple behavioral changes that are stably expressed for minutes after repeated aversive stimulation. Specifically, animals increase their speed and reduce their defecation and feeding rates. In addition, after the first 2-3 stimuli, animals scale these behavioral responses with further incoming aversive stimuli, showing a form of sensitization. Our work here extends a previous description of the speeding response^34^. We focused on optogenetic ASH activation and chemical activation by copper, but ASH also responds to harsh mechanical stimuli that may be processed differently by downstream circuits^55^. In addition, the aversive state that we examined here is reminiscent of an aversive state in which repeated cross-modal stimulation induced an RID-dependent increase in speed^36^. Finally, we have previously characterized a stable aversive response to a single noxious thermal stimulus, which reduces feeding and causes repeated turning^41^. This latter state involved ADA activation, but not AVH activation, suggesting some degree of overlap. Overall, there are a variety of distinct aversive states expressed by *C. elegans* that manifest as stable responses to specific aversive cues. It is possible that combinatorial coding by a set of integrator neurons capable of functioning in parallel may confer this specificity.

Where does the long timescale stability come from in the neural integrator circuits studied here? These behavioral changes occur in a backdrop of serotonergic modulation. All the animals studied here were exposed to dense bacterial food, which acts through serotonin to drive stable dwelling states^18,49,56–59^. The aversive stimuli change multiple behavioral parameters that regress back to the dwelling baseline over minutes. We found that the neural integrators that elicit these changes utilize multiple distinct stability mechanisms. Stable activation of PVQ can be evoked by direct PVQ activation in synaptic transmission mutants, suggesting a cell-intrinsic mechanism for persistent activity. In contrast, this does not work for AVH stimulation, which require upstream ASH stimulation for stable activation. Moreover, AVH signaling via the neuropeptide NLP-10 is required for stable expression of the behavioral state, which may extend the timeframe of the behavioral effects beyond that of AVH’s persistent activity. Overall, this suggests that cell-intrinsic factors, circuit interactions, and neuromodulatory signaling are all involved in generating this aversive behavioral state. Future mechanistic studies could utilize this experimental system to dissect the complex interplay between these factors in controlling long timescale behaviors.

## Acknowledgments

We thank Evan Ardiel, Anne Hart, Linlin Fan, and members of the Flavell lab for critical reading of the manuscript. We thank Eviatar Yemini, Oliver Hobert, Cori Bargmann, and CGC for strains and plasmids. S.N.B. acknowledges funding from a Picower Postdoctoral Fellowship. S.W.F. acknowledges funding from NIH (DC020484, GM135413); The Brain Research Foundation; The Picower Institute for Learning and Memory; the Howard Hughes Medical Institute; and The Freedom Together Foundation. S.W.F. is an investigator of the Howard Hughes Medical Institute.

## Author Contributions

Conceptualization, S. N. B., A.K., S.P., J.K., K.M., S.W.F. Methodology, S. N. B., A.K., S.P., J.K., K.M., S.W.F. Software, S. N. B., A.K., S.P., C.K., A. H., J.K., S.W.F. Formal analysis, S.N.B., A.K., A.H. Investigation, S.N.B., A.K., S.P., F.M., C.E., D.K., E.B. Writing – Original Draft, S.N.B. and S.W.F. Writing – Review & Editing, S.N.B. and S.W.F. Funding Acquisition, S. N. B., S.W.F.

## Methods

### Strain List

| Genotype | Source | Strain Name |
| --- | --- | --- |
| Wild-type Bristol N2 | Caenorhabditis Genetics Center (CGC) | N2 |
| *egl-21(n476)* IV | CGC | KP2018 |
| *sbt-1(ok901)* V | CGC | RB987 |
| *flp-9(ok2730)* IV | CGC | RB2067 |
| *F15A4.5(gk3259)* II; *flp-5(gk3123)* X | CGC | VC3280 |
| *flp-26(gk3015)* X | CGC | VC3017 |
| *flp-16(ok3085)* II | CGC | RB2275 |
| *flp-5(tm12132)* X | National BioResource Project (NBRP) | FX31781 |
| *nlp-8(syb762)* IV | CGC | HBR2317 |
| *nlp-15(ok1512)* I | CGC | VC1063 |
| *nlp-9(sy1451)* V | CGC | PS8676 |
| *nlp-21(sy1807)* III | CGC | PS9520 |
| *nlp-19(sy1457)* X | CGC | PS8682 |
| *nlp-17(ok3461)* IV | CGC | RB2498 |
| *nlp-9(tm3572)* V | NBRP | FX03572 |
| *nlp-14(tm1880)* X | NBRP | FX01880 |
| *nlp-21(tm2569)* III | NBRP | FX02569 |
| *nlp-36(tm5156)* III | NBRP | FX05156 |
| *nlp-80(sy1264)* V | CGC | PS8307 |
| *nlp-10(tm6332)* III | NBRP | FX06232 |
| *nlp-11(syb530)* II | CGC | PHX530 |
| *nlp-50(sy2074)* II | CGC | PS10162 |
| *flvEx317[nlp-76::CoChR, nlp-76::mScarlett, myo-3::mCherry]* | this study | SWF716 |
| *flvEx493[nlp-76::Chrimson; nlp-76::wrmScarlet; nlp-17::GCaMP7f; nlp-17::GCaMP7f; flvEx400[Pnlp-76::GCaMP6m (60 ng/ul), Pnlp-76::mScarlet (20 ng/ul)]* | this study | SWF1246 |
| *flvEx710[Pgpa-6::CoChR, myo-2::mCherry]* | this study | SWF1403 |
| *flvEx710[Pgpa-6::CoChR, myo-2::mCherry]; flvEx548[nlp-76::Chrimson (15ng/ul), myo-3::mCherry]* | this study | SWF1404 |
| *flvEx548[nlp-76::crimson (15ng/ul), myo-3::mCherry]* | this study | SWF1098 |
| *flvEx141[Prig-3::CochR (50ng/ul), Pmyo-2::mCherry]* | this study | SWF382 |
| *flvEx489[Psto-3::CoChR (50 ng/ul), Psto-3::mScarlet (10ng/ul), Pmyo-3::mCherry (5ng/ul)]* | this study | SWF1007 |
| *flvEx366[Pinx-1::CoChR (50ng/ul); Pmyo-3::mCherry (5ng/ul)]* | this study | SWF782 |
| *flvEx448[glr-1::Cre (18ng/uL); tdc-1::inv(CoChR)::sl2GFP 30ng/uL); myo-2::mCherry (5ng/uL)]* | Kramer et al., 2026 | SWF946 |
| *lite-1(ce314); gur-3(ok2245); otIs670[NeuroPAL]; flvIs17[tag-168::NLS-GCaMP7F, gcy-28.d::NLStag-* | Atanas et al., 2023 | SWF702 |
| *RFPT, ceh-36::NLS-tag-RFPT, inx-1::tag-RFPT, mod-1::tagRFPT, tph1(short)::NLS-tag-RFPT, gcy-5::NLS-tag-RFPT, gcy-7::NLStag-RFPT]* |  |  |
| *flvEx377[sra-6::GCaMP7f (30 ng/ul), sra-6::wormScarlett (10 ng/ul)]* | this study | SWF802 |
| *flvEx619[nlp-76::Chrimson; nlp-76::wormScarlett; nlp-82::GCaMP7f, elt-2::mCherry]* | this study | SWF1256 |
| *flvEx604[nlp-76::Chrimson; nlp-76::wormScarlett; h1h-34::GCaMP7f; elt-2::mCherry]* | this study | SWF1218 |
| *flvEx542[nlp-76::TeTx::mCherry, myo-2::mCherry]* | this study | SWF1091 |
| *flvEx317[nlp-76::CoChR, nlp-76::mScarlett, myo-3::mCherry]; flvEx540[eat-4::nCre (5ng/ul), flp-26::inv(unc-103(gf), GFP) (30ng/ul), myo-2::mCherry]* | this study | SWF961 |
| *flvEx317[nlp-76::CoChR, nlp-76::mScarlett, myo-3::mCherry]; flvEx582[Phlh-34::unc-103::SL2::GFP, Pelt-2::mCherry]* | this study | SWF1189 |
| *flvEx317[nlp-76::CoChR, nlp-76::mScarlett, myo-3::mCherry]; flvEx268[nlp-17::unc-103(gf) (50ng/ul); myo-2::mCherry]* | this study | SWF807 |
| *flvEx488[Peat-4::nCre (10ng/ul), Pflp-26::inv(CoChR,GFP) (30ng/ul), Pmyo-3::mCherry (5ng/ul)]* | this study | SWF1006 |
| *flvEx592[Phlh-34::CoChR::SL2::GFP, Pmyo-3::mCherry]* | this study | SWF1175 |
| *flvEx498[Pnlp-17::CoChR (30 ng/ul), Pmyo-3::mCherry(5 ng/ul)]* | this study | SWF1019 |
| *flvEx548[nlp-76::Chrimson (15ng/ul), myo-3::mCherry]; flvEx578[Pvab-7::nCre, Pflp-26::inv(CoChR::SL2::GFP), Pmyo-2::mCherry]* | this study | SWF1207 |
| *flvEx578[Pvab-7::nCre, Pflp-26::inv(CoChR::SL2::GFP), Pmyo-2::mCherry]* | this study | SWF1149 |
| *flvEx548[nlp-76::Chrimson (15ng/ul), myo-3::mCherry]; flvEx580[Phlh-34::CoChR::SL2::GFP, Pmyo-2::mCherry]* | this study | SWF1209 |
| *flvEx580[Phlh-34::CoChR::SL2::GFP, Pmyo-2::mCherry]* | this study | SWF1151 |
| *flvEx548[nlp-76::Chrimson (15ng/ul), myo-3::mCherry]; flvEx144[Pnlp-17::CoChR (30ng/ul),Pmyo-2::mCherry]* | this study | SWF1208 |
| *flvEx144[Pnlp-17::CoChR (30ng/ul),Pmyo-2::mCherry]* | this study | SWF385 |
| *flvEx488[Peat-4::nCre (10ng/ul), Pflp-26::inv(CoChR,GFP) (30ng/ul), Pmyo-* | this study | SWF1090 |
| 3::mCherry (5ng/ul)]; flvEx268[nlp-17::unc-103(gf) (50ng/ul); myo-2::mCherry] |  |  |
| flvEx592[Phlh-34::CoChR::SL2::GFP, Pmyo-3::mCherry]; flvEx540[eat-4::nCre (5ng/ul), flp-26::inv(unc-103(gf), GFP) (30ng/ul), myo-2::mCherry] | this study | SWF1177 |
| flvEx592[Phlh-34::CoChR::SL2::GFP, Pmyo-3::mCherry]; flvEx268[nlp-17::unc-103(gf) (50ng/ul); myo-2::mCherry] | this study | SWF1176 |
| flvEx498[Pnlp-17::CoChR (30 ng/ul), Pmyo-3::mCherry(5 ng/ul)]; flvEx540[eat-4::nCre (5ng/ul), flp-26::inv(unc-103(gf), GFP) (30ng/ul)] | this study | SWF1082 |
| nlp-10(tm6232); flvEx317[nlp-76::CoChR, nlp-76::mScarlett, myo-3::mCherry] | this study | SWF958 |
| nlp-10(tm6232); flvEx592[Phlh-34::CoChR::SL2::GFP, Pmyo-3::mCherry] | this study | SWF1234 |
| nlp-10(tm6232); flvEx498[Pnlp-17::CoChR (30 ng/ul), Pmyo-3::mCherry(5 ng/ul)] | this study | SWF957 |
| nlp-10(tm6232); flvEx317[nlp-76::CoChR, nlp-76::mScarlett, myo-3::mCherry]; flvEx610[Phlh-34::nlp-10cDNA, Pmyo-2::mCherry] | this study | SWF1235 |
| flvEx789[hhlh-34::Chrimson, hhlh-34::GCaMP7f, myo-3::mCherry] | this study | SWF1564 |
| unc-13(s69) I; flvEx789[hhlh-34::Chrimson, hhlh-34::GCaMP7f, myo-3::mCherry] | this study | SWF1567 |
| flvEx583[Pnlp-17::Chrimson, Pnlp17::GCaMP7f, Pmyo3::mCherry] | this study | SWF1154 |
| unc-13(s69) I; flvEx583[Pnlp-17::Chrimson, Pnlp17::GCaMP7f, Pmyo3::mCherry] | this study | SWF1155 |
| flvEx583[Pnlp-17::Chrimson, Pnlp17::GCaMP7f, Pmyo3::mCherry]; flvEx379[nlp-17::TeTx (50 ng/ul), elt-2::nGFP (5 ng/ul)] | this study | SWF1321 |
| flvEx317[nlp-76::CoChR, nlp-76::mScarlett, myo-3::mCherry]; eat-4(ky1091, ky1092) | this study | SWF1311 |
| flvEx317[nlp-76::CoChR, nlp-76::mScarlett, myo-3::mCherry]; eat-4(ky1091, ky1092); flvEx650[Pnlp-17::FLP, Pmyo-2::mCherry] | this study | SWF1314 |
| flvEx498[Pnlp-17::CoChR (30 ng/ul), Pmyo-3::mCherry(5 ng/ul)]; eat-4(ky1091, ky1092); flvEx650[Pnlp-17::FLP, Pmyo-2::mCherry] | this study | SWF1312 |

### C. elegans genetics and growth conditions

*C. elegans* Bristol strain N2 was used as wild type. Animals were maintained on NGM agar plates seeded with *E. coli* OP50 bacteria and kept at 22°C, 40% humidity. One day-old adults were used for all experiments. For genetic crosses, genotypes were confirmed by PCR and/or sequencing. Transgenic animals were generated by injecting DNA with fluorescent co-injection markers into the gonads of young adult hermaphrodites.

### Plasmids, Promoters, and Transgenics

Plasmid backbones: The plasmids containing GCaMP6m^60^, NLS-GCaMP7f^41^, Cre^61^, Flp^47^, unc-103(gf)^62^, TeTx^61^, Chrimson^59^ or CoChR^62^ open reading frames in the pSM vector backbone have been previously described. The intersectional promoter backbone, consisting of the inverted/floxed vector was previously described^18^. For *nlp-10* rescue, we used the *nlp-10* cDNA inserted into the pSM vector backbone for expression.

Promoters used in this study: *nlp-76*, *flp-26*, *nlp-82, nlp-17*, *hlh-34*, *sra-6*, *sto-3*, *gpa-6*, *eat-4*, *gcy-13*, *inx-1*, *rig-3*.

The conditional *eat-4* knockout allele was derived from a previously described CRISPR/Cas9 genome edited strain (CX17641)^47^, in which the endogenous *eat-4* locus was flanked by FRT sites with mCherry inserted after the stop codon. PVQ-specific Flp expression under the *nlp-17* promoter led to *eat-4* excision and mCherry expression specifically in PVQ neurons, confirming both that there is bona fide *eat-4* expression in PVQ and the functionality of the cell-specific knockout.

### Multi-animal behavioral recordings

Multi-animal recordings of *C. elegans* locomotion were conducted as previously described^59^. For optogenetic experiments, L4-stage animals were picked onto plates with bacterial lawns containing all-trans-retinal (ATR: 0.5uM and 50uM ATR for dual-color vs single-color optogenetic experiments, respectively) 16-20 hours before the experiment. On the day of the recordings, one day old adult animals were transferred to 10cm NGM plates seeded with OP50 bacteria, placed on the recording stage, and allowed to acclimatize for 15 minutes. All animals were recorded using Streampix software at 3 fps. JAI SP-20000M-USB3 CMOS cameras (5120×3840, mono) with Nikon Micro-NIKKOR 55mm f/2.8 were used. IR-panel LEDs (EFFILUX) provided backlighting. Videos were analyzed using previously-described custom MATLAB scripts^63^. For optogenetic stimulation, light was supplied from a 470nm (for CoChR; 3, 5 and 10 uW/mm^2^ for Extended Data Figure 1; 0.2 uW/mm^2^ for dual-color optogenetics, and 10 uW/mm^2^ for all other optogenetics experiments with blue light), or 625nm (for Chrimson; 50 uW/mm^2^ for dual color optogenetics) Mightex LED at defined times in the video. For dual-color optogenetics experiments, additional optical filters (Thorlabs 450 nm shortpass filter; Chroma ET630/20x filter) were used to clean up the LED light and spectrally separate the red and blue excitation wavelengths. Change in speed was measured from the 1-minute period preceding the first optogenetic stimulus, unless indicated otherwise. Half-decay time following the last stimulus was estimated by bootstrapped resampling of the total variation-denoised mean speed trace. The half-decay point was defined as the first time point at which smoothed speed was sustained below the midpoint between the peak post-stimulus speed and pre-stimulus baseline, with standard deviation reported across iterations.

### Behavioral modeling from optogenetic stimuli

To predict speed traces from optogenetic stimuli, we used training data from recordings with optogenetic ASH stimulation at interstimulus intervals of 4, 16, and 64 seconds. The model was validated on 32-second interval data and tested on datasets with 8-second interstimulus intervals. We compared a family of stimulus-response models that only differed in how they integrate past stimuli over time. The same splits of training, validation, and test data were used to train all models. In all models, predicted speed was computed as a gain-and-offset transformation of historical and present optogenetic stimuli, where stimulus sensitivity (c_s) scaled the response magnitude and baseline speed (b) set resting locomotion speed in the absence of stimulation. The tested kernel families were: (i) a leaky integrator with exponential decay governed by decay rate τ (exponential decay), (ii) a finite-memory boxcar kernel with uniform weights over window length W (rectangular), (iii) a recency-weighted finite-memory kernel with linearly decreasing weights over window length W (linear decay), and (iv) a delayed response kernel peaked at lag μ with temporal spread σ (Gaussian). We also included a memoryless baseline model that responded only to instantaneous stimulus (linear, no-decay model).

For model training, we normalized all speed traces to the 99th percentile of the maximum speed observed in the training dataset. We then obtained 100 bootstrap samples (with replacement) of the speed traces in the training data. Models were fit by minimizing the mean squared error (MSE) on the bootstrapped samples, using the JAX (jax.value_and_grad for automatic differentiation), Equinox (eqx.apply_updates for model class definition and updates) and Optax (optax.adam for gradient-based optimization with Adam optimizer) Python packages. Model convergence was determined based on the loss on withheld validation data (which was distinct from the testing data, as described in the main text). For each model, we fit the shared parameters (c_s, b) along with the kernel-specific parameters and evaluated predictive performance on the 8-second interstimulus test condition. For the exponential decay model, we further examined the MSE of the fits after parameter degradation on an independent dataset collected on different days with various optogenetic stimulus paradigms (5+5 grouped stimulus dataset).

### Single-animal behavioral recordings

For joint recordings of pumping, defecation and locomotion, we used previously described custom-built single worm tracking microscopes^64^. These custom microscopes have a live-tracking function that permits long-term recording of single moving animals at 20 frames per second. L4s animals were picked 16-20 hours before the recording day onto 50uM ATR plates. On the day of the recording, animals were transferred to low-peptone (0.4g/1L) NGM plates seeded with OP50 the day before the recording (thin bacterial lawns are required to track freely-moving animals on the microscopes). LabView software controlled the microscope and acquired the images. Data were then analyzed in R Studio and MATLAB. For optogenetic stimulation, 532nm laser light was supplied at defined times at an intensity of 123 uW/mm^2^. Prior to analysis, speed and pumping rates were smoothed using median filtering with a 1-second and 10-second window, respectively. Defecation events were smoothed using a 1-minute moving mean.

### Copper boundary encounters

To present animals with an aversive stimulus during imaging, a round section of flat agar NGM containing 2 mM copper chloride was excised using an 8 mm diameter biopsy punch. Hot regular NGM was then poured into the resulting hole and allowed to cool, plugging it with neutral agar. The pad was subsequently flipped, creating a circular neutral arena in the center surrounded by a sharp copper gradient. Both agar layers were sandwiched beneath a glass microscope slide to ensure uniform thickness. Next, 10 µL of liquid NGM buffer containing 20x concentrated OP50 was placed on top of the agar pad. For whole brain imaging, 4 µL of microsphere beads suspended in liquid NGM were pipetted on the outer edges of the agar pad, away from the central arena, to reduce coverslip pressure on the animal. Encounters with the copper gradient were scored by measuring the distance between each animal’s head and the copper boundary. Reversals initiated within 1.125mm of this boundary were counted as copper-evoked reversals. Each copper-evoked or spontaneous reversal was categorized as an isolated versus repeated reversal according to the number of same-type reversals initiated within the previous 30 seconds (0, or one or more).

### Freely-moving calcium imaging in individual neurons

Calcium imaging of individual neurons in freely-moving animals was conducted as previously described with a few small modifications^65^. During copper boundary encounters, animals were mounted on flat agar surrounded by a round agar boundary with 2 mM copper along with 20x concentrated OP50 bacterial food (see methods on Copper boundary encounter for more details). They were covered with cover glass and GCaMP/mCherry data was recorded at 5 fps with 10ms exposure. Pulse illumination from an X-cite LED source was used for excitation, along with a Chroma 59022x multi-band bandpass filter. For optogenetics experiments, animals were transferred onto 0.25uM ATR plates the day before the experiment. After the 16-20 hours, animals were mounted onto flat agar pads with 20x OP50 food and covered in cover glass. The GCaMP fluorescence was recorded at 10 fps with 10ms exposure. Excitation was again from an X-cite LED light source, but a Chroma 49002 filter set was used for these experiments. For optogenetic stimulation, 625nm LED light (Mightex) filtered through Chroma ET630/20x filter and was supplied at defined times at an intensity of 200 uW/mm^2^.

Imaging was performed with a 4x/0.2NA objective and data was acquired on two Andor Zyla 4.2 Plus sCMOS cameras or Teledyne Photometrics Prime BSI sCMOS cameras. A Cairn TwinCam beam splitter was used to separate GCaMP and mCherry signals. The GCaMP/mCherry imaging at 5 fps was interleaved with 5 fps brightfield imaging, achieved via NI-DAQ triggering of different light sources using the NIS Elements Illumination Sequence module. For optogenetics experiments combined with GCaMP imaging, GCaMP images at 10 fps was interleaved with 10 fps brightfield imaging.

For data analysis, the neuronal soma was tracked (using the bright mCherry signal for ratiometric imaging; or GCaMP fluorescence during optogenetics experiments with Chrimson) using custom ImageJ macros. After cell positions were determined by ImageJ tracking, the GCaMP and mCherry signals were extracted from each ROI at each time point. Background was subtracted from each signal and then the ratio of the two background-subtracted fluorescence measurements was taken. For optogenetics experiments, GCaMP fluorescence was normalized to 30 seconds pre-stimulus period by subtracting the average baseline signal from this each subsequent timepoint.

### Whole-brain imaging

Whole-brain imaging in moving animals was conducted as previously described^41^. Recordings were performed using the transgenic strain SWF702, which expresses pan-neuronal GCaMP and NeuroPAL, and carries *lite-1* and *gur-3* null mutations. To present animals with an aversive stimulus during imaging, a round section of flat agar NGM containing 2 mM copper chloride was excised using an 8 mm diameter biopsy punch. Hot regular NGM was then poured into the resulting hole and allowed to cool, plugging it with neutral agar. The pad was subsequently flipped, creating a circular neutral arena in the center surrounded by a sharp copper gradient. Both agar layers were sandwiched beneath a glass coverslip to ensure uniform thickness. Next, 10 µL of liquid NGM buffer containing 20x concentrated OP50 was placed on the central arena of the agar pad, with 4 µL of microsphere beads suspended in liquid NGM positioned away from the central arena, to reduce coverslip pressure on the animal. Day 1 adults were mounted on the central NGM arena, covered with cover glass and imaged for 16 minutes. Fluorescence data and behavioral traces were extracted from videos as previously described^41,42^. For whole-brain imaging behavioral quantification, reversals were defined as periods of backward velocity.

### Wet copper drop response assay

To measure responses to copper, we used 2mM copper chloride dissolved in M13 buffer, as previously described. The day before the assay, 10 cm assay plates were seeded with 2 ml of OP50, and lids were left open for 2–3 hours in a biosafety hood to allow the bacterial lawn to dry. L4 larvae were picked onto OP50-seeded NGM plates and grown for 16-20 hours to obtain day 1 adults. On the day of the experiment, adult animals were transferred to seeded assay plates (5 animals per plate) and allowed to acclimate for 30 minutes. Each animal was then presented with a liquid copper droplet as it moved forward. Animals that paused or reversed within 4 seconds were scored as responders. For reversing animals, the number of body bends during reversal was counted and normalized to N2 wild-type controls recorded on the same day. All experiments were performed blind to genotype.

### Statistics and Reproducibility

The statistical tests used in the study are described in the accompanying figure legends. Non-parametric statistical tests were used and multiple comparison corrections were applied, as described in the figure legends.

## Data availability

Data are freely and publicly available at https://doi.org/10.5061/dryad.w9ghx3g4v.

## Code availability

Code for analysis of whole-brain calcium imaging data has been previously described^41^ and is available at https://github.com/flavell-lab/ANTSUN.

---

**Extended Data Figure 1.**
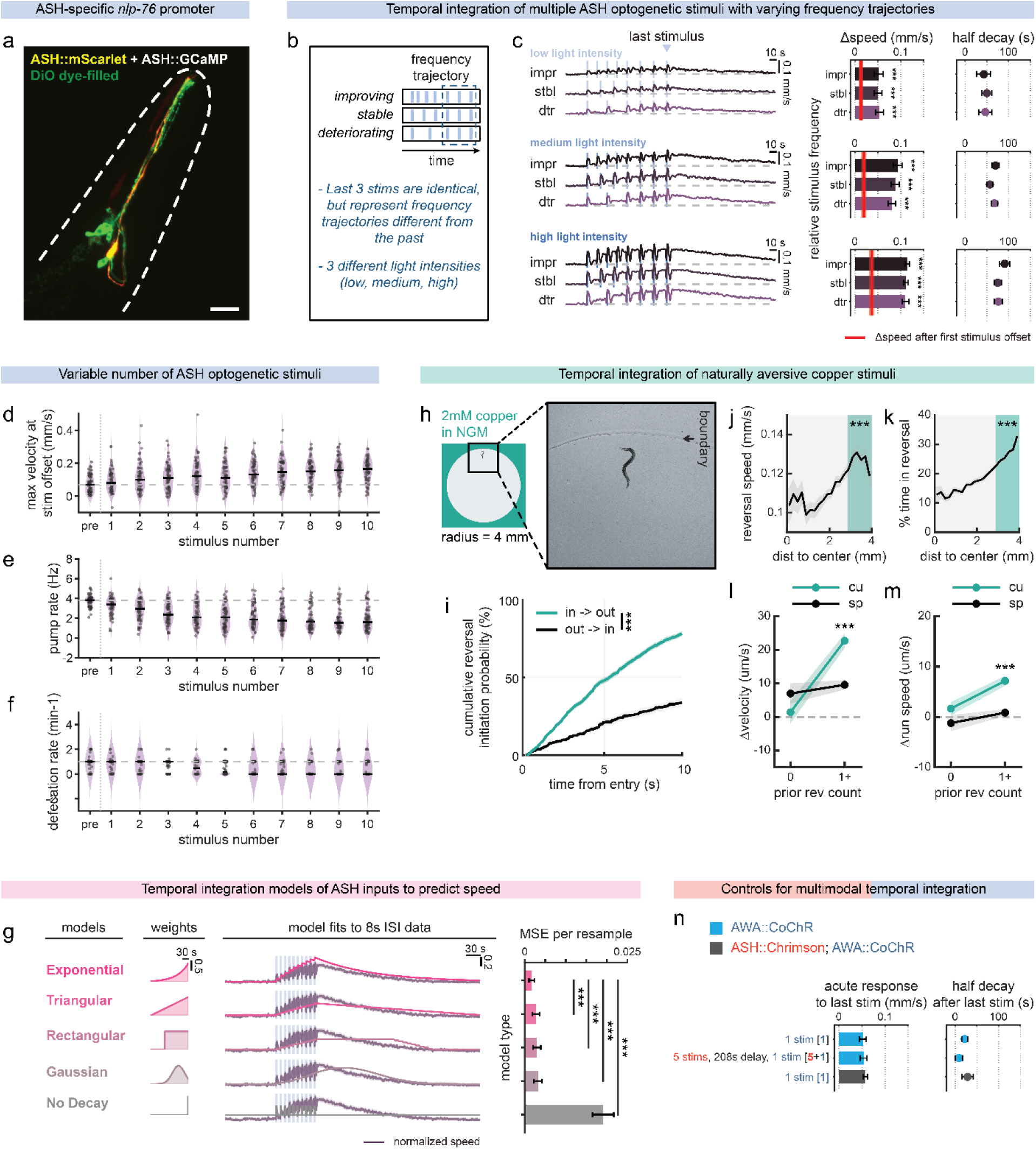
a. ASH specific nlp-76 promoter driving mScarlet expression in an animal dye-filled with DiO. b. Schematic showing optogenetic ASH stimulation patterns. These patterns hold constant the most recent optogenetic stimuli, while varying the earlier stimuli. In the “improving” conditions, the stimuli gradually decrease in frequency during the stimulus pattern, whereas in the “deteriorating” conditions, the stimuli gradually increase in frequency. c. Change in locomotion speed during optogenetic ASH stimulation at varying relative frequencies and intensities, based on stimulus patterns schematized in (b). The experiment was run at 3 different light intensities (low, medium and high). There was no difference based on “improving”, “stable”, or “deteriorating” at any light intensity. d. Maximum velocity at stimulus offsets among individual ASH::CoChR animals after receiving varying numbers of light stimuli (1x to 10x 2-second stimuli at 8 second intervals). Same animals as shown in Figure 1b. Note that the entire distribution gradually shifts upwards with increasing number of stimuli, rather than, for example, a bi-modal distribution emerging with increasing number of stimuli. e. Average pumping rate with repeated ASH stimuli. Same animas are shown as in Figure 1c. f. Average defecation rate with repeated ASH stimuli. Same animas are shown as in Figure 1d. g. Models and fits for predicting locomotion speed based on ASH optogenetic stimuli as inputs. Second column: shape of learned temporal kernels for each model family, shown as a function of lag (seconds before the current time). Third column: example predictions; the measured mean normalized speed is shown in purple (shaded region, ± SEM), and the model prediction is overlaid in pink hues. Blue shading indicates ASH optogenetic stimuli. Each model predicts speed as a gain-and-offset transformation of the stimulus trace with the corresponding kernel weights. Model families: Exponential (exponentially decaying “leaky integrator”), Rectangular Block (finite window), Triangular Block (finite window with linearly decreasing recency weights), and Gaussian Block (peaked kernel centered at a lag). Text annotations report the mean-squared error (MSE) for each model across bootstrapped resamples. Right: average MSE across bootstrap resamples for each model family (error bars, ± s.d. across resamples) on withheld testing data. Asterisks indicate significance of pairwise comparisons to the exponential model using an empirical resampling-based two-sided test. h. Illustration of encounters with the copper boundary. Left: schematic. Right: representative image of a worm in arena. Note the visible boundary (arrows). i. Cumulative probability of reversal initiation over time, comparing entries into aversive (in->out) versus neutral (out->in) zones. Shows only the reversals initiated after crossing on forward runs. Lines show the average across 77 animals. Shaded bands indicate SEM. Area under the cumulative distribution functions were compared using a paired Wilcoxon signed-rank test. ***p<0.001. j. Reversal speeds as it varies with distance to copper boundary. 0 indicates the center, while a distance of 4mm is directly on the boundary. Shaded bands indicate SEM across animals. Reversals that were initiated in the teal-shaded region were categorized as copper-evoked reversals and compared to spontaneous reversals from central arena. Paired Wilcoxon signed-rank test. ***p<0.001. k. Percent time spent in reversals with varying distances to the copper boundary, normalized to the time each animal spent in that bin. Shaded band indicates SEM across animals. Analyzed the same way as in j. l. Change in velocity post- versus pre-reversal for copper-evoked (CU) and spontaneous (SP) reversals, with different numbers of past reversals of the same type over the last 30 seconds. Velocity was averaged over 30 s before and after reversal. To isolate velocity changes attributable to the reversal itself, trials were excluded for both CU and SP if the animal subsequently reapproached the copper boundary or initiated another copper-evoked reversal within 30 s. Error bars show SEM. Wilcoxon rank-sum test. ***p<0.001. m. Change in forward run velocity post- versus pre-reversal for copper-evoked (CU) and spontaneous (SP) reversals, with different numbers of past reversals of the same type over the last 30 seconds. Analyzed the same way as in l. n. AWA::CoChR controls for dual-color optogenetics setup. Left: magnitude of sensory responses to the AWA stimulus, measured from the 8-second baseline period preceding it. Right: persistence of the speeding response. All conditions compared to AWA::CoChR controls receiving a single AWA stimulus with no prior optogenetic stimulation. For additional ASH::Chrimson controls, see Extended Data Figure 4.

**Extended Data Figure 2.**
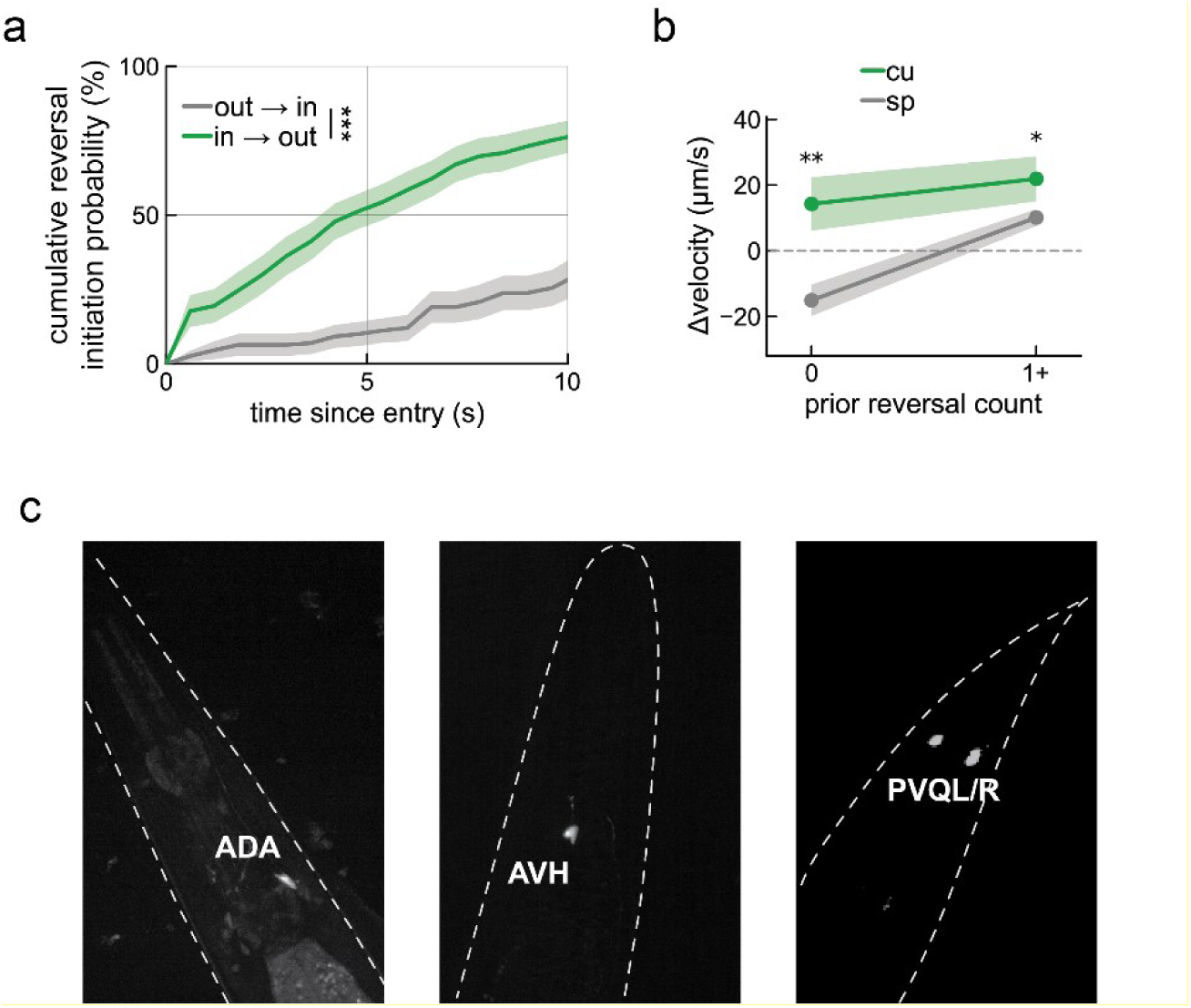
a. In whole-brain imaging animals, cumulative probability of reversal initiation over time, comparing entries into aversive (in->out), copper-containing area versus exits from the outer copper area into the central neutral (out->in) zones. Shows only the reversals initiated after crossing on forward runs. Lines show the average across 27 animals. Shaded bands indicate SEM. Area under the cumulative distribution functions were compared using a paired Wilcoxon signed-rank test. ***p<0.001. b. Change in velocity post- versus pre-reversal for copper-evoked (CU) and spontaneous (SP) reversals, with different numbers of past reversals of the same type over the last 30 seconds. Velocity was averaged over 30 s before and after reversal. To isolate velocity changes attributable to the reversal itself, trials were excluded for both CU and SP if the animal subsequently reapproached the copper boundary or initiated another copper-evoked reversal within 30 s. Error bars show SEM. Wilcoxon rank-sum test. ***p<0.001. c. Promoters used to study ADA (*Pnlp-82::inv(loxP::unc-103(gf)::SL2::GFP::loxP); Peat-4::nCre*), AVH (*Phlh-34::GCaMP7f*) and PVQ (*Pnlp-17::GCaMP7f*).

**Extended Data Figure 3.**
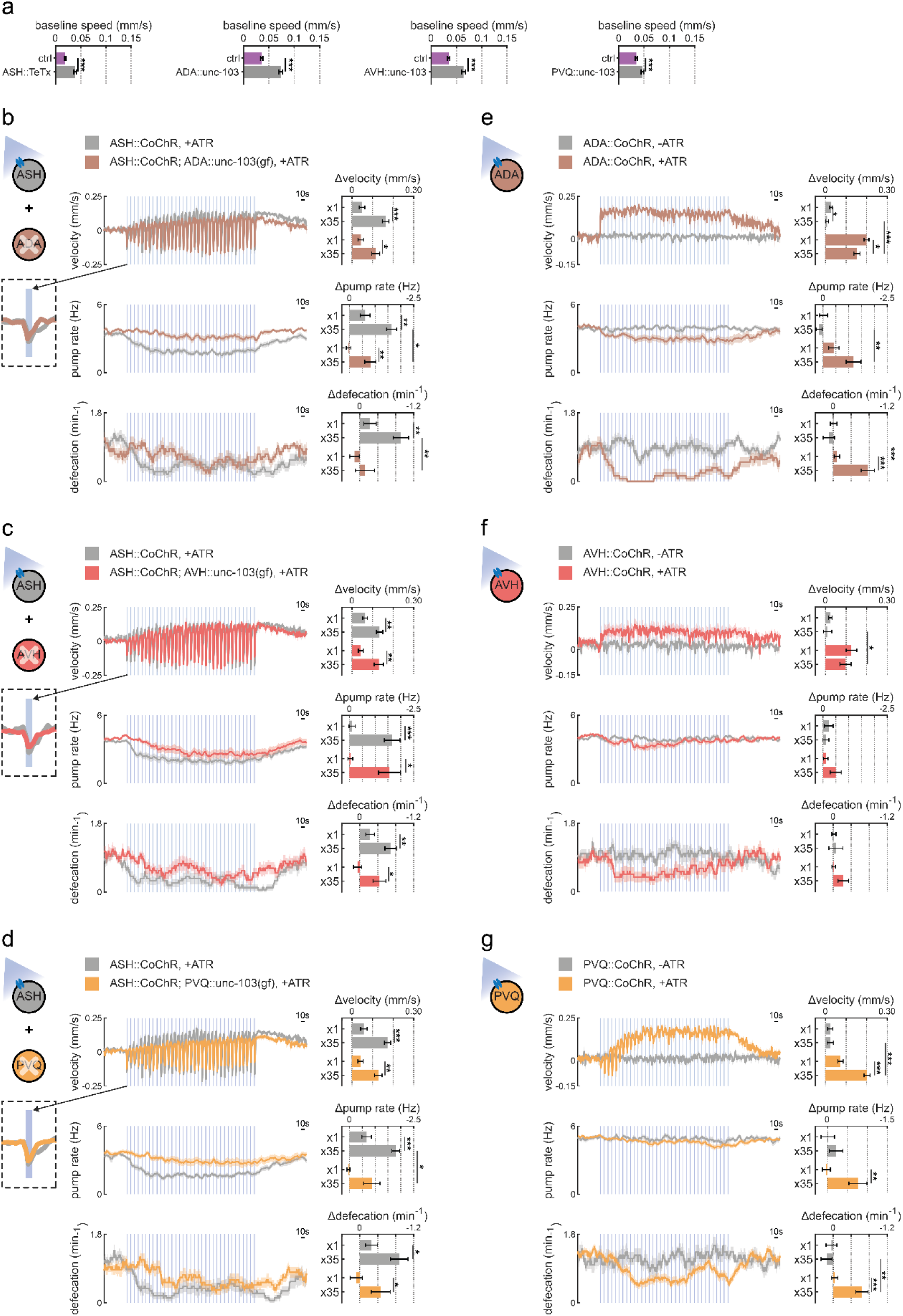
a. Baseline speed (prior to any optogenetic stimulation) in ASH::CoChR animals also expressing ASH::TeTx, ADA::unc-103gf, AVH::unc-103gf and PVQ::unc103gf animals. “ctrl” is animals with only the ASH::CoChR transgene. ***p<0.001, Wilcoxon rank-sum test. Note that there is a mild increase in baseline speed in the neural perturbation lines, suggesting that there is not a generic motor deficit in these animals. b. Schematic illustrating optogenetic activation of ASH in animals with silenced ADA (ADA::unc-103gf). Event-triggered changes in ADA-silenced animals for velocity (top), pumping rate (middle) and defecation rate (bottom) after repeated ASH optogenetic activation (left), and comparisons of changes after single vs repeated ASH stimulation (right). Insets show velocity during the first stimulus response. Each behavior was compared to day-matched wild type controls. *p<0.05, ***p<0.001, Wilcoxon rank-sum test c. Schematic illustrating optogenetic activation of ASH in animals with silenced AVH (AVH::unc-103gf), shown as in (b). d. Schematic illustrating optogenetic activation of ASH in animals with silenced PVQ (PVQ::unc-103gf), shown as in (b). e. Schematic illustrating behavioral changes during optogenetic activation of the ADA neurons. Event-triggered changes in velocity (top), pumping rate (middle) and defecation frequency (bottom) after repeated ADA optogenetic activation (left), and comparisons of changes after single vs repeated ADA stimulation (right). Each behavior was compared to day-matched no ATR controls. *p<0.05, ***p<0.001, Wilcoxon rank-sum test. f. Schematic illustrating optogenetic activation of the AVH neurons, shown as in (f). g. Schematic illustrating optogenetic activation of the PVQ neurons, shown as in (f).

**Extended Data Figure 4.**
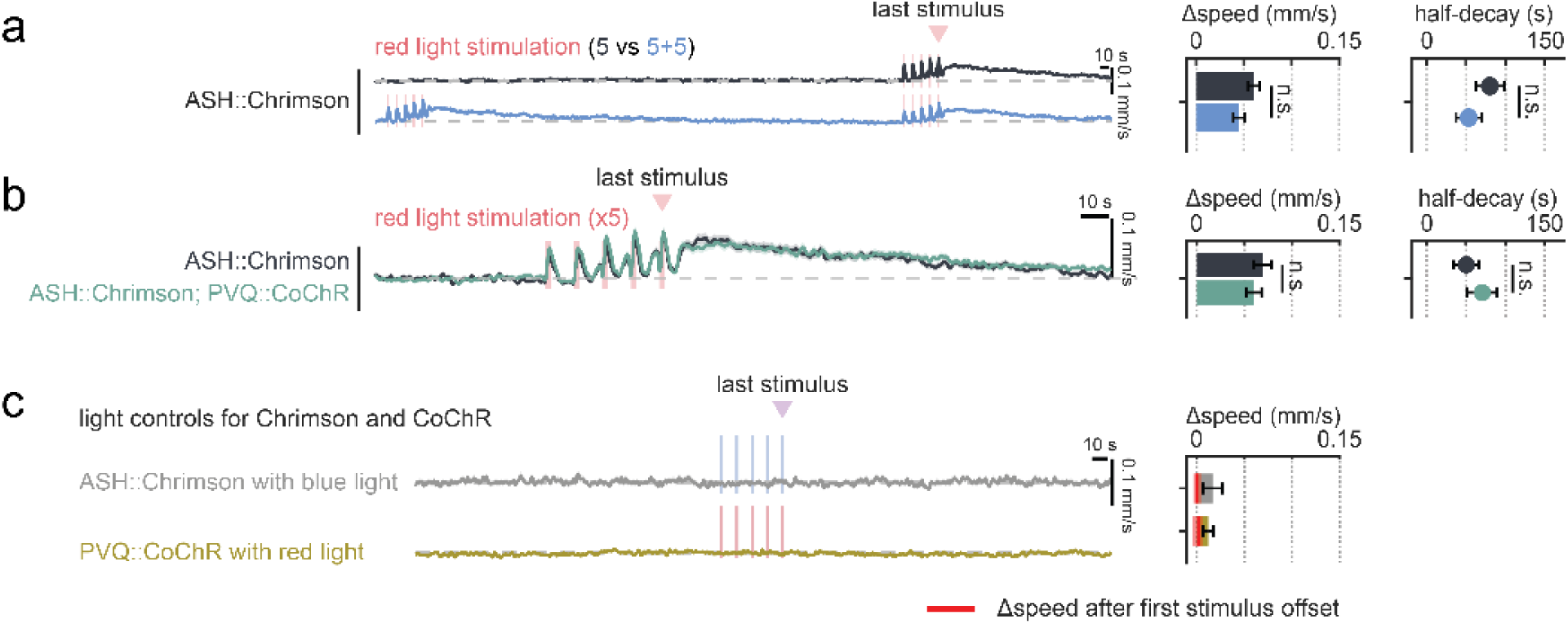
a. Responses in ASH::Chrimson animals when they were stimulated 5x, or twice in groups of 5 with 558 seconds of delay. b. Responses to 5x red light stimuli in ASH::Chrimson transgenic animals versus ASH::Chrimson;PVQ::CoChR animals. Note that the responses are the same in these two strains when there was no prior blue activation. c. Light controls for Chrimson and CoChR, showing ASH::Chrimson and PVQ::CoChR animals stimulated with blue and red light, respectively. Note that red light fails to induce behavior in CoChR animals and blue light fails to induce behavior in Chrimson animals.

**Extended Data Figure 5.**
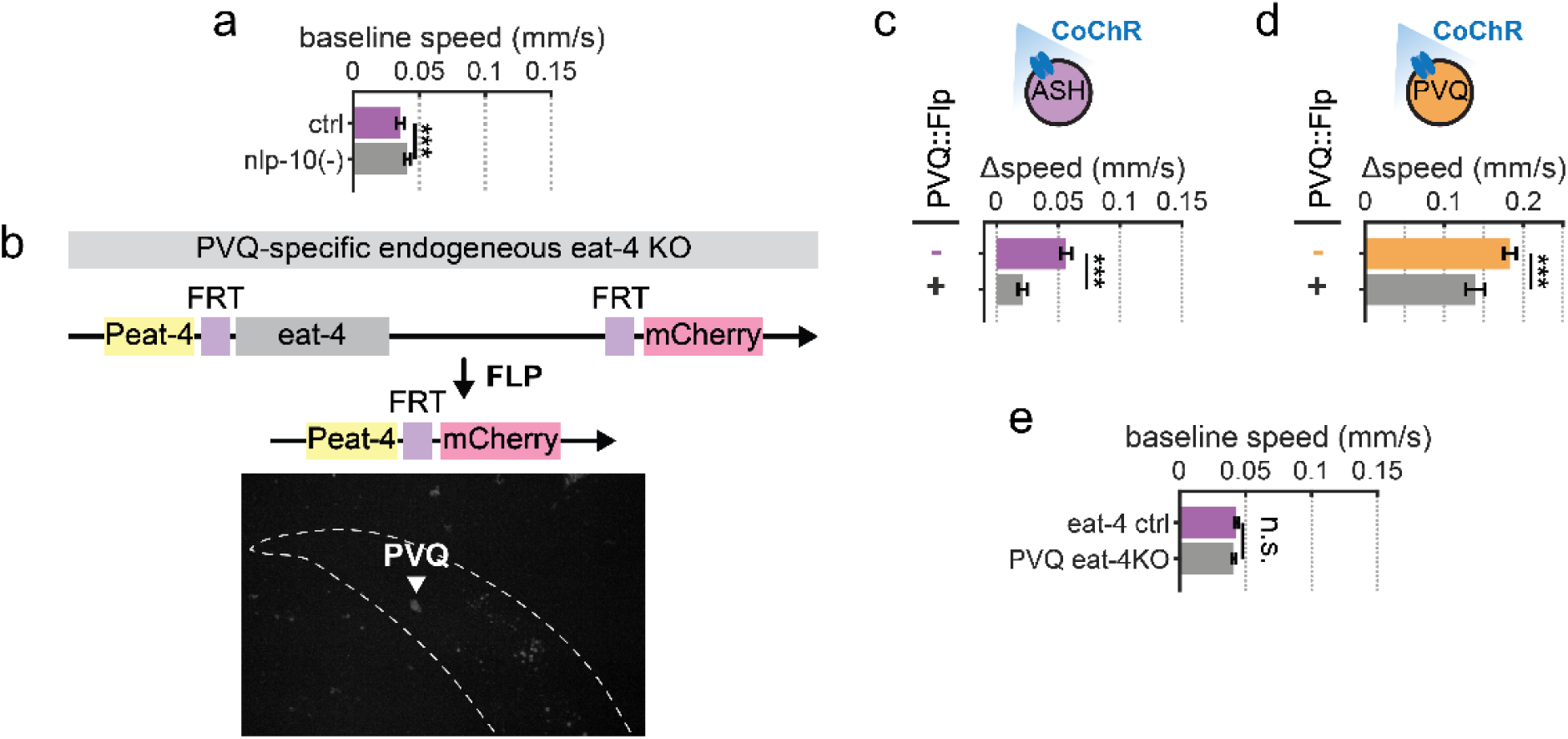
a. Baseline speed of ASH::CoChR animals with *nlp-10* null mutation, compared to ctrl animals with ASH::CoChR in an otherwise wild-type background. ***p<0.001, Wilcoxon rank-sum test. b. Top: Schematic of transgenes used for PVQ neuron specific eat-4 knockout (based on plasmids from^47^). Bottom: representative image of PVQ mCherry expression in the resulting strain. c. Behavioral effects of optogenetic ASH activation in *FRT::eat-4::FRT* control animals vs. animals with PVQ-specific deletion of *eat-4* to disrupt glutamatergic signaling. Cell-specific deletion was achieved using the *eat-4* conditional deletion strain^47^. ***p<0.001, empirical p-value. d. Behavioral effects of optogenetic PVQ activation in control animals vs. animals with PVQ-specific deletion of eat-4 to disrupt glutamatergic signaling. ***p<0.001, empirical p-value. e. Baseline speed in animals with PVQ-specific deletion of *eat-4*, compared to *FRT::eat-4::FRT* control animals.

